# Interacting active surfaces: a model for three-dimensional cell aggregates

**DOI:** 10.1101/2022.03.21.484343

**Authors:** Alejandro Torres-Sánchez, Max Kerr Winter, Guillaume Salbreux

## Abstract

We introduce a modelling and simulation framework for cell aggregates in three dimensions based on interacting active surfaces. Cell mechanics is captured by a physical description of the acto-myosin cortex that includes cortical flows, viscous forces, active tensions, and bending moments. Cells interact with each other via short-range forces capturing the effect of adhesion molecules. We discretise the model constitutive equations using a finite element method, and provide a parallel implementation in C++. We discuss examples of application of this framework to simulations involving small and medium-sized aggregates: we consider the shape and dynamics of a cell doublet, a planar cell sheet, and a growing cell aggregate. This framework opens the door to the systematic exploration of the cell to tissue-scale mechanics of cell aggregates, which plays a key role in the morphogenesis of embryos and organoids.

**Author summary:** Understanding how tissue-scale morphogenesis arises from cell mechanics and cell-cell interactions is a fundamental question in developmental biology. Here we propose a mathematical and numerical framework to address this question. In this framework, each cell is described as an active surface representing the cell acto-myosin cortex, subjected to flows and shape changes according to active tensions, and to interaction with neighbouring cells in the tissue. Our method describes cellular processes such as cortical flows, cell adhesion, and cell shape changes in a deforming three-dimensional cellular aggregate. To solve the equations numerically, we employ a finite element discretisation, which allows us to solve for flows and cell shape changes with arbitrary resolution. We discuss applications of our framework to describe cell-cell adhesion in doublets, three-dimensional cell shape in a simple epithelium, and three-dimensional growth of a cell aggregate.

## 1 Introduction

Tissue morphogenesis relies on the controlled generation of the cellular forces that collectively drive tissue-scale flows and deformation [1,2]. The interplay between cell-cell adhesion, cellular mechanics and the cytoskeleton plays a key role in determining how biological tissues self-organise [3]. These ingredients are also crucial for the growth of *in vitro* organoids, organ-like structures derived from stem cells which can self-organise into complex structures reminiscent of actual organs [4–6].

Several classes of models have been proposed to describe the mechanics of multicellular aggregates, such as cellular Potts models [7–13], phase field models [14–18] and vertex models [19]. Vertex models, and the closely related Voronoi vertex models [20,21], describe cells in a tissue as polyhedra that share faces, edges and vertices forming a three-dimensional junctional network [19,22–24]. Cell deformations are encoded by the displacement of the vertices ***X***_*a*_ of the network. These displacements are dictated by vertex forces ***F***_*a*_ stemming from cell pressures *P_c_*, surface tensions *t_f_*, and line tensions Γ_*e*_ that are coupled to virtual changes in cell volume *δV_c_*, face area *δA_f_*, and edge length *δl_e_* respectively in a work function

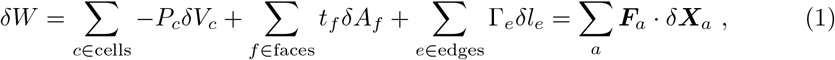

where to get to the last expression one needs to express *δV_c_*, *δA_f_* and *δl_e_* in terms of a virtual displacement of the vertices *δ****X***_*a*_. Two and three-dimensional versions of the vertex model have been employed for many applications, for instance to study cell packing [23,25], cell sorting [26], wound closure [27], cyst formation [28], tumourigenesis in tubular epithelia [29] among many other [19]. However because of their definition, vertex models do not explicitly resolve cortical flows on the cell surface. The effect of cell-cell adhesion is also implicitly introduced in the surface tension *t_f_*, which mix together physical processes arising from molecular bonds between cells and surface forces arising in the cell membrane and in the actomyosin cytoskeleton. In vertex models with vertices positions as degrees of freedom, topological transitions leading to cell-neighbour exchange are encoded explicitly by formulating rules to change edges in the network.

At the single cell scale, a number of studies have shown the relevance of coarse-grained, continuum models to describe the mechanics of the cell surface. In this approach, an active fluid theory taking into account cellular cortical flows, gradients of active cytoskeletal tension and their regulation, and orientation and filament alignment in the actin cortex, has proven successful to describe the mechanics of cell polarisation, cell motility or cell division [30–39]. From a computational perspective, there has been a growing attention to the simulation of the dynamics of fluid interfaces both with prescribed [40–43] and with time-evolving shape [38, 39, 44–50]. However, to our knowledge no computational framework has attempted to provide a physical description of three-dimensional cellular aggregates taking into account explicitly the mechanics of a single cell surface described as an active surface, as well as cell-cell adhesions.

Here we bridge this gap and introduce a new modelling and simulation framework, and a freely available code [51], for the mechanics of cell aggregates in three dimensions (Fig. 1). We describe cells as interacting active surfaces [52]. The governing equations for the cell surface mechanics are discretised using a finite element method. In this method, each cell is represented by a three-dimensional mesh with vertices positions ***X***_*a*_. In analogy with Eq. (1) in vertex models, we start from the virtual work theorem for interfaces (Eq. (2)) and find the net forces at the vertices ***F***_*a*_ (Eq. (11)), which vanish in the absence of inertia. This condition imposes cortical flows and cell shape changes. In this framework, topological transitions appear as a natural output of the remodelling of cell-cell interactions and are not treated explicitely.

**Fig 1.**
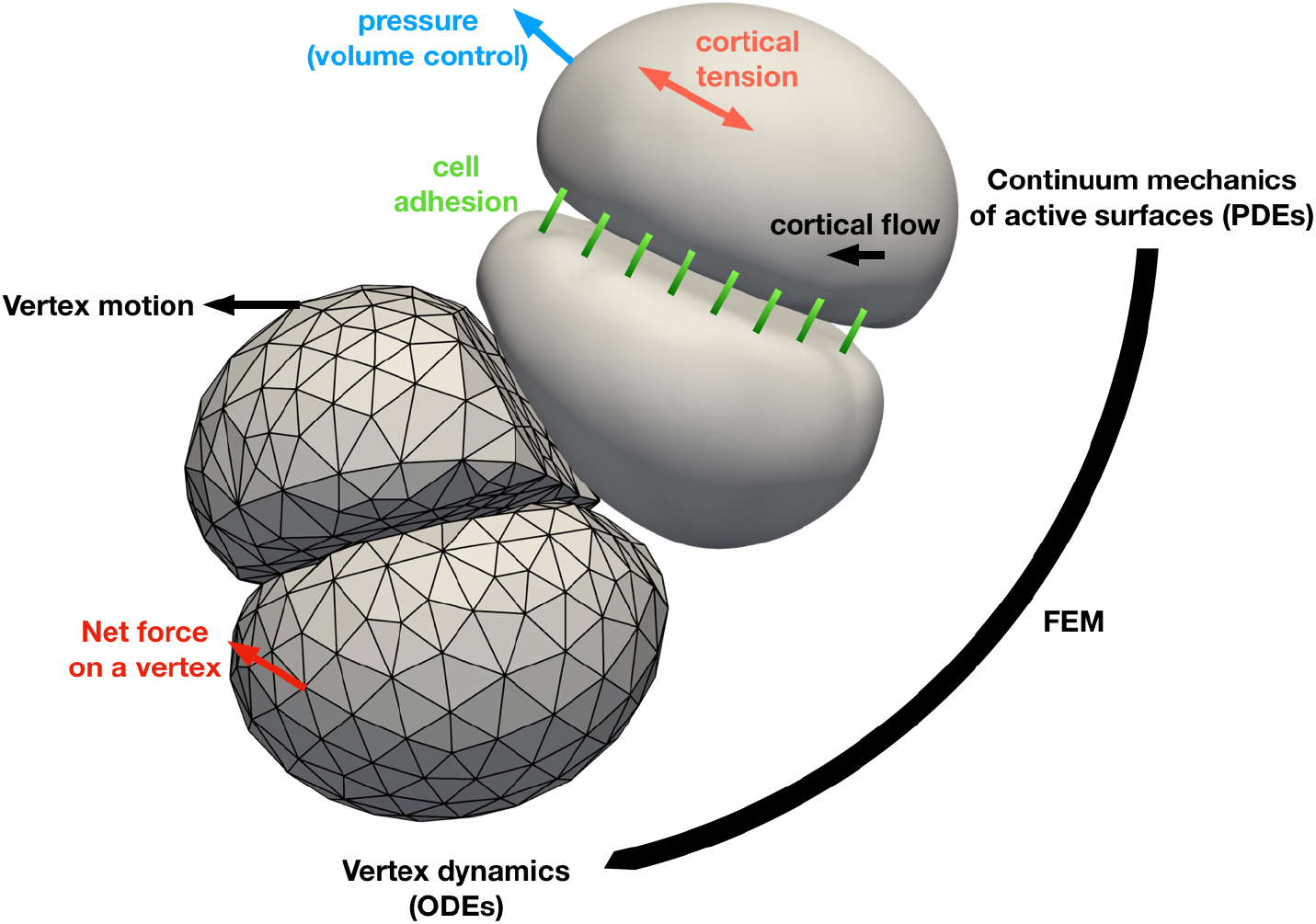
Schematic of the interacting active surface framework. A tissue is described by a collection of triangular meshes representing each cell. The dynamics of the tissue is described by the dynamics of the vertices making the cellular meshes, similar to how the movement of the vertices of a vertex model describe the deformation of a tissue. The motion of the cell mesh is obtained by coarse-graining continuum mechanics equations of a theory of active surfaces via the finite element method. In this theory, cortical flows, cortical tensions, intracellular pressures, bending moments and forces arising from cell-cell interactions are taken into account.

The main benefits of our method are that it (1) can incorporate complex descriptions of the physics of the cell surface, including sources of tension, in-plane and normal moments [52], (2) accounts for cell-cell adhesion explicitly through constitutive laws that can be adapted to represent different biological scenarios, (3) resolves cell shape with arbitrary resolution given by the mesh size of the finite element discretisation.

The nonlinear equations of the model are treated computationally with the use of nonlinear solvers. Since we consider only the discretisation of the cell surface, the number of degrees of freedom is considerably smaller than in 3D Cellular Potts or phase-field models, both of which require a 3D discretisation. On the other hand, to capture cell shape, cortical flows and cell-cell interactions accurately, we need to use many more vertices per cell than in a vertex model, which leads to a greater computational cost. To limit computational time, the code is parallelised to allow each cell to be stored on a different partition, each possibly using a pool of cores (hybrid MPI-OpenMP method). As such, it is possible to simulate several tens of cells on a computing cluster.

We now turn to the description of our framework. In Section 2 we describe the mathematical formulation and the discretisation of our method. We show some examples of application in Section 3 and end in Section 4 with conclusions, summary and ideas for future work.

## 2 Materials and methods

In this section we describe the main elements of our framework. We start by introducing the governing equations for a single, isolated cell described as a fluid active surface. We use a virtual work formulation for the mechanics of a surface, together with a finite element discretisation, to obtain the equations that dictate the movement of the vertices of the mesh. We introduce cell-cell interactions, represented by an interaction potential between pairs of surfaces, that result in a force density and a tension acting on each cell. Finally, we describe a mesh reparametrisation method that allows simulations to handle large tangential deformations of the surface which can arise for continuously flowing fluid interfaces such as the cell cortex.

### 2.1 Governing equations and discretisation for a single surface

#### 2.1.1 Virtual work for interfaces

Our starting point is the statement of virtual work for a closed interface 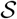 representing a single cell. We describe 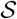 with a parametrisation ***X*** (*s*^1^, *s*^2^) where *s*^1^, *s*^2^ are two surface coordinates and ***X*** a point in the 3D space in which the surface is embedded. We denote the coordinates of the 3D cartesian basis by greek indices *α, β*,…, and the coordinates on the surface by latin indices *i,j*…. Here and elsewhere in the manuscript we use Einstein summation convention for repeated indices. Given the tangent vectors ***e***_*i*_ = *∂*_*i*_***X*** one can compute the metric tensor *g_ij_* = ***e***_*i*_ · ***e***_*j*_ and the curvature tensor *C_ij_* = – ***n*** · *∂_i_****e***_*j*_, where ***n*** = (***e***_1_ × ***e***_2_)/|***e***_1_ × ***e***_2_| is the outer normal to the surface. Other notations of differential geometry are given in S1 Appendix 1. Here and in the following, we consider the limit of low Reynolds number where inertial terms are negligible, a limit relevant to the physics at the cell scale which is of interest here [53]. The mechanics of a single surface can then be described by the following statement of the principle of virtual work for 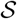 [52]:

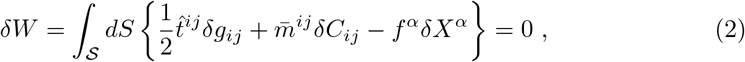

where *δ**X*** is an infinitesimal displacement of the surface, and *δg_ij_* and *δC_ij_* the associated infinitesimal variation of the metric and curvature tensors. Expressions for *δg_ij_* and *δC_ij_* in terms of *δX^α^* are given in S1 Appendix 2. Eq. (2) relates infinitesimal variations of geometric quantities of the interface to their work conjugates: the external force density *f^α^* coupled to the surface displacement, the tension tensor 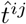 related to metric variations, and the bending moment tensor 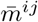 coupled to variations of the curvature tensor. Eq. (2) is equivalent to the statement of balance of linear and angular momentum at low Reynolds number (S1 Appendix 3). We have neglected external and internal normal moments for simplicity, which lead to extra terms in Eq. (2) (S1 Appendix 3).

#### 2.1.2 Constitutive laws

To describe the mechanics of a single surface, we need to specify constitutive equations for the mechanical tensors 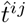 and 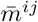. We denote by ***v*** the velocity field on the cell surface. Here, we assume that the cell surface can be represented as an active viscous layer, with a bending rigidity and a spontaneous curvature. As a result, we use the following constitutive equations:

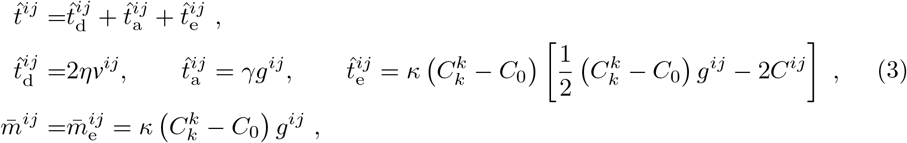

where *v_ij_* is the strain-rate tensor on the surface:

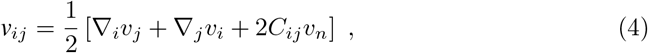

with ∇_*i*_ the covariant derivative operator defined in S1 Appendix 1. Here, *η* is the surface viscosity; for simplicity we do not explicitly distinguish between the shear and bulk viscosity of the surface. *γ* is the surface tension of the cell, which is not necessarily homogeneous on the cell surface. As we expect this contribution to largely arise from active forces generated in the actomyosin cortex [54], we later refer to it as the active tension. The contributions 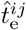 and 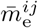 arise from an effective Helfrich free energy penalising the membrane curvature with bending modulus *κ* and spontaneous curvature *C*_0_ (S1 Appendix 4).

The external force density acting on a single cell is split into two contributions

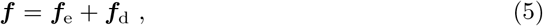

arising respectively from the action of the pressure difference across the surface

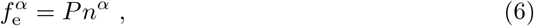

and from an effective external friction force density

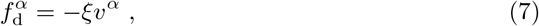

where *ξ* is a friction coefficient. In the following, the pressure *P* is adjusted to impose a value of the cell volume. Eq. (2), together with the constitutive equations (3)-(7), leads to a complete set of equations to determine the velocity field ***v*** on the cell surface.

The constitutive laws Eqs. (3)-(7) are a simple choice of physical description of the surface. Other terms, for instance additional viscous or active bending moments, could be introduced: linear irreversible thermodynamics provides with a set of additional possible terms that could play a role in the dynamics of isotropic, polar or nematic active surfaces and could be added to the framework described here [52, 55].

#### 2.1.3 Finite element discretisation

We now discretise the geometry of the surface using finite elements based on a triangular control mesh of *N_e_* triangles or elements and *N_v_* vertices (Fig. 2). The parametrisation of 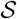 is of the form

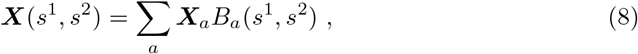

where the sum is taken over vertices of the control mesh, labelled by *a*, ***X***_*a*_ is the position of vertex *a* on the control mesh, and 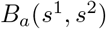 is its basis function, defined in terms of coordinates (*s*^1^, *s*^2^) on the control mesh. Given that 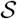 may have bending or in-plane moments coupled to variations of the curvature tensor and the Christoffel symbols in the differential virtual work (see Eq. (2) and Eq. (34) in S1 Appendix 3), the basis functions *B_a_* must be chosen to have second order derivatives that are square-integrable; here we follow [38,56,57] and use Loop subdivision surfaces, which lead to smooth surfaces that satisfy this condition by construction. We note that, in practice, a consistent parametrisation (*s*^1^, *s*^2^) of the entire control mesh is not needed: indeed one can consider the surface 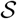 as a union of surface elements associated to each triangle e of the control mesh, which are given in terms of barycentric coordinates 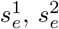, of the element *e* by

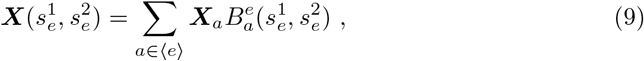

see Fig. 2. Here we have denoted by 〈*e*〉 the set of vertices whose basis functions are nonzero in element *e*, and 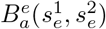 corresponds to the contribution of the basis function *B_a_*, within element *e*, parametrised by the barycentric coordinates of *e*. For Loop subdivision surfaces, 〈*e*〉 is formed by the vertices in the triangular element *e*, and all first neighbours of these vertices.

**Fig 2.**
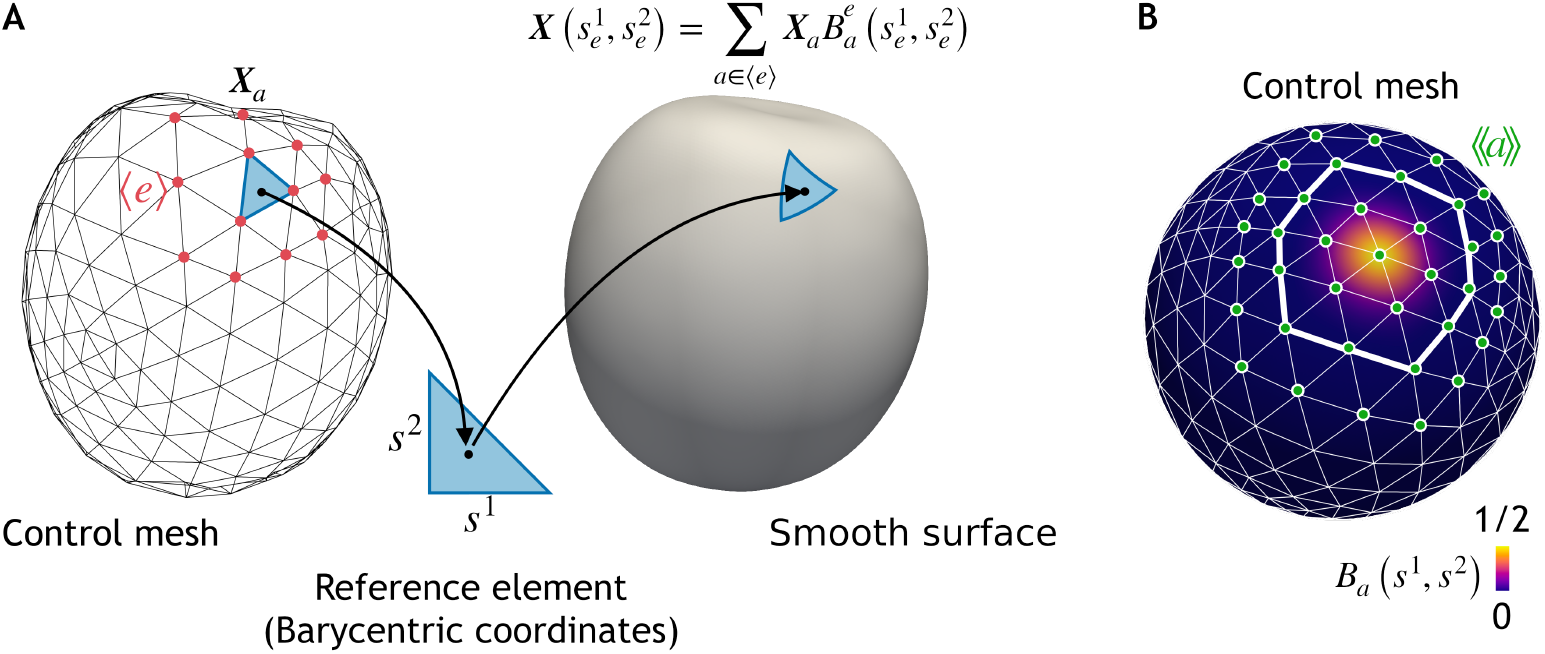
A smooth surface 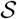 representing a cell is obtained based on a triangular control mesh with vertex positions ***X***_*a*_ (A) and a set of basis functions per vertex *a* (B). (A) To define the mapping between the control mesh and the cell surface, the barycentric coordinates of points in a triangular element *e* in the control mesh 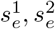, which span a reference triangle, are used to define a point on the cell surface 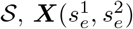 (Eq. (9)). Points 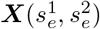 are obtained by summing basis functions 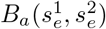, weighted by ***X***_*a*_, over vertices a whose basis functions have a non-zero contribution to this element, an ensemble denoted 〈*e*〉. (B) Example of the basis function associated to a vertex in the mesh. For Loop subdivision surfaces basis functions, the basis function spans the first and second rows of elements surrounding the vertex (thicker white line). The vertices that interact with vertex *a* in the same cell, represented by the set 〈〈*a*〉〉 (green) are formed by the first, second, and third nearest neighbours in the mesh.

The discretisation of the surface (Eq. (8)) transforms the virtual work principle (Eq. (2)) into an expression of the form

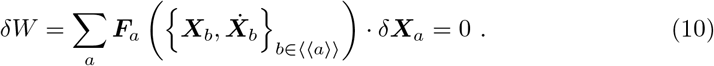

Here ***F***_*a*_, which can be interpreted as the net force on vertex *a*, can be obtained by substituting, in the differential virtual work, the analytical expressions for the variations *δg_ij_* and *δC_ij_* in terms of *δ***X**** = ∑_*a*_ *B_a_*(*s*^1^, *s*^2^)*δ****X***_*a*_ (S1 Appendix 6). We have denoted by 〈〈*a*〉〉 the set of vertices interacting with vertex *a*, which for Loop subdivision surfaces is formed by its first, second and third ring of neighbours, see Fig. 2B. Since Eq. (10) has to be satisfied for any *δ****X***_*a*_, the net forces on the vertices need to vanish

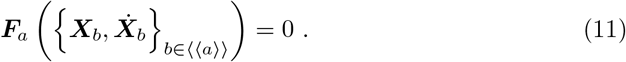

Through this discretisation, we transform the original continuum problem into a set of coupled ordinary differential equations (ODEs). These ODEs need to be discretised in time to be resolved computationally; in the following we denote by (*n*) the *n*–th time step of the time evolution. Here we employ a semi-implicit Euler discretisation, where terms arising from the effective bending energy and active tension are discretised in a fully implicit manner, whereas viscous and frictional terms are treated explicitly. This particular choice leads to a variational time-integrator that preserves the dissipative structure of the dynamics for a homogeneous and time-independent active tension [38]. This implies that the tension and bending moment tensors at step *n* are evaluated as:

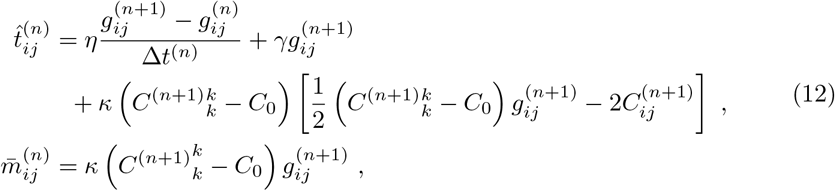

which can be written in terms of 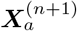 and 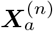 through the relation between the surface definition and the position of vertices, Eq. (8). To discretise the strain rate tensor, we have used its relation with the rate of change of components of the metric tensor (see S1 Appendix 2). Plugging these expressions in the differential virtual work Eq. (2) allows us to transform Eq. (11) into a set of (nonlinear) algebraic equations (S1 Appendix 6)

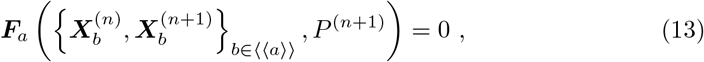

which can be solved using a Newton-Raphson method together with the discretisation of the volume constraint, which is imposed through the nonlinear constraint

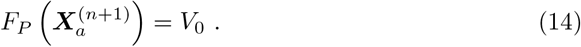

Here *P*^(*n*+1)^ is the intracellular pressure, playing the role of a Lagrange multiplier imposing conservation of volume, see S1 Appendix 6.2. In this method, the solution is updated by 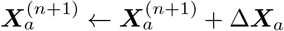, *P*^(*n*+1)^ ← *P*^(*n*+1)^ + Δ*P*^(*n*+1)^ where Δ***X***_*a*_ and Δ*P*^(*n*+1)^ satisfy the linearised equations

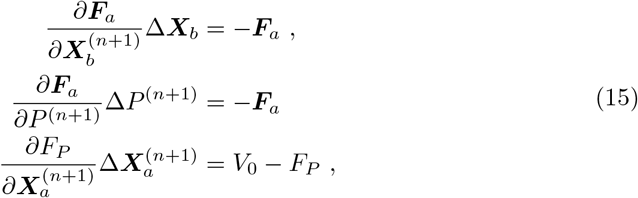

until the norm of ***F***_*a*_ and *F_P_* – *V*_0_ is below a given tolerance. Here 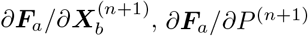, and 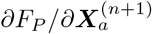 form the tangent matrix. The tangent matrix is symmetric, in particular 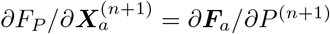. Because only vertices in 〈〈*a*〉〉 interact with *a,* this matrix is sparse and the linear system can be solved efficiently with an iterative solver. Our computational framework makes use of Trilinos [58] to handle all linear algebra objects, including sparse matrices, in parallel.

### 2.2 Forces arising from cell-cell interactions

Cell-cell adhesion modifies the previous equations by introducing interactions between the vertices of interacting surfaces. We now consider a set of surfaces 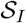, *I* =1… *N* describing *N* interacting cells. We assume that a pair of distinct cells *I, J* interact via an effective energy:

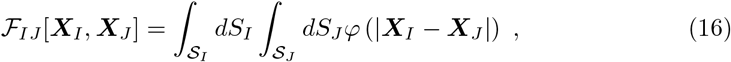

where *φ* is an effective adhesion potential. We discuss below a microscopic motivation for this potential. The virtual work differential of the cell aggregate can then be written as:

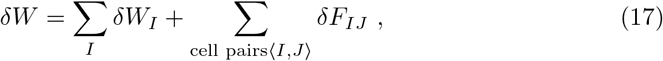

where the contribution *δW_I_* is given by Eq. (2) with infinitesimal displacement (*δ****X***_*I*_), metric variation (*δg_I_ij__*) and curvature variation (*δC_I_ij__*) of cell *I*, and with the tension tensor 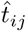, bending moment tensor 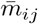, and external force density excluding cell-cell interaction forces ***f***, given by the constitutive equations (3)-(7). Here and in the following, we denote distinct cell pairs 〈*I, J*〉, such that in sums taken over 〈*I, J*〉, each pair is counted only once.

Variations of the interaction free energy lead to

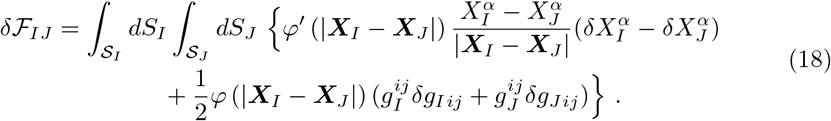

Comparing this expression with Eq. (2), each cell interaction with a cell *J* is contributing an additional external force density on cell *I*, ***f***_*I J*_, and an additional isotropic tension to cell *I*, 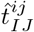:

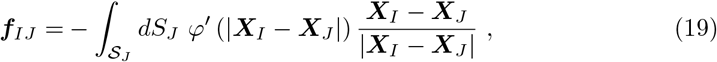

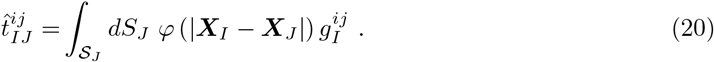

In addition, although this point is not directly apparent from Eq. (18), the variation 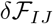 can be written only in terms of normal displacements, provided that *φ* only depends on |***X***_*I*_ – ***X***_*J*_| (Eq. (53) in S1 Appendix 3). This shows that the net driving force from the interaction potential, including the effect of both the force density ***f***_*I J*_ and tension 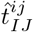, has a vanishing tangential component. Intuitively, the interaction energy does not change if cells do not change shape, so it cannot generate a driving force for tangential motion.

Following the same procedure as in the previous section, the condition *δW* = 0, with *δW* defined in Eq. (17), gives rise to a set of (nonlinear) algebraic equations of the form

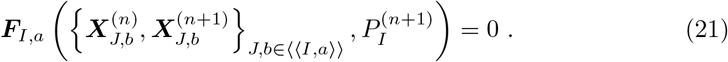

Here 〈〈*I, a*〉〉 identifies the set of vertices (identified as the pair of labels *J* for the cell considered and *b* for the vertex considered) that interact with vertex *a* in cell *I*.

The form of the effective energy (Eq. (16)) can be motivated microscopically by considering an ensemble of stretchable linkers connecting pairs of surfaces, which quickly equilibrate by binding and unbinding to cell surfaces, and whose free concentration is set by contact with a reservoir. We characterise such an ensemble by a two-point concentration field *c_IJ_* (***X***_*I*_, ***X***_*J*_), which quantifies the number of bound linkers joining the points ***X***_*I*_ and ***X***_*J*_ per unit area of the first and second surfaces (thus, it has units of the inverse of an area squared). The concentration *c_I_* denotes the concentration of unbound, free linkers in cell *I*, with units of the inverse of an area. In the dilute limit, the free energy of this ensemble can be written as:

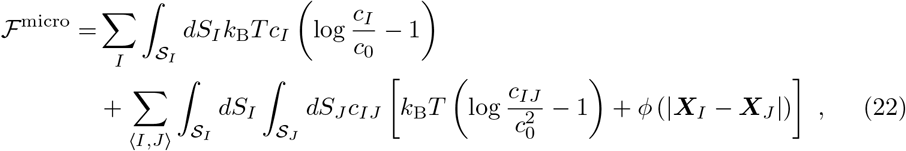

where *k*_B_ is the Boltzmann constant and *T* the temperature. The first sum corresponds to the free energy of free linkers, described as an ideal solution in contact with a reservoir imposing a chemical potential. This chemical potential determines the value of *c*_0_. The second sum corresponds to the free energy of bound linkers, described as an ideal solution, with an energy per linker dependent on the linker elongation, quantified by the potential *ϕ*. The potential is defined such that the force sustained by a linker of length *r* is –*ϕ*′(*r*). One can then show that if linkers are at equilibrium with respect to surfaces *I, J* with a fixed shape, the free energy of interaction of two surfaces *I, J* is (S1 Appendix 5)

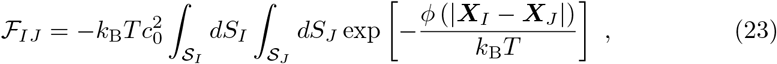

which gives a relation between the microscopic behaviour of the linkers and the potential *φ* introduced in Eq. (16). For the particular choice of *ϕ*(*r*) = *ϕ*_0_ + *k*(*r* – *r*_min_)^2^/2 with *k* the bond stiffness and *r*_min_ its reference length, one obtains:

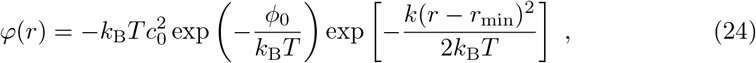

which corresponds to an inverted Gaussian with centre at the equilibrium length *r*_min_, width 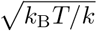 and depth 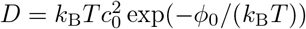.

This description however still does not take into account short range repulsion between two cells surfaces. This could be taken into account by introducing a second repulsive interaction potential between surfaces. Here we choose instead to introduce a convenient effective potential of interaction, the Morse potential

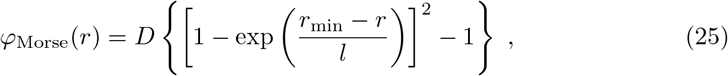

which like the interaction potential in Eq. (23), vanishes for *r* → ∞, has a minimum at *r* = *r*_min_ with minimum value –*D*, and is also peaked around its minimum with characteristic length *l*. In addition, for *l* ≪ *r*_min_ it exhibits a sharp short-range repulsion.

Although *φ*_Morse_(*r*) decays with *r* rapidly, it is convenient to have a strict cut-off on its range, to limit interacting vertices of the meshes. Therefore, we further multiply *φ*_Morse_(*r*) by a smooth step function:

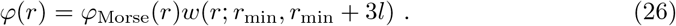

where

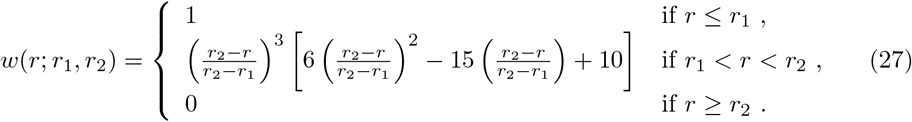

This ensures that the potential *φ*(*r*) goes to zero exactly at a distance *r*_min_ + 3*l* with first and second order continuous derivatives; a convenient property to solve numerically the non-linear equations (Eq. (21)).

### 2.3 Surface reparametrisation

The method discussed in the previous sections is based on a Lagrangian scheme, such that a node of the mesh flows with the material particles of the interface. This, however, can lead to large in-plane distortions, notably if surface tension gradients generate in-plane flows [38, 47]. To compensate for the resulting mesh distortion, we describe here a reparametrisation method, in the spirit of Refs. [56, 57]. In this method, vertices of the control mesh move tangentially to the surface to minimise an effective mesh quality energy. We stress that this step does not bear any physical meaning. Here we discuss again a single cell. Given the mesh of a cell, we define the energy

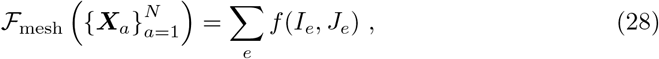

where the sum is performed over the triangles of the control mesh, and

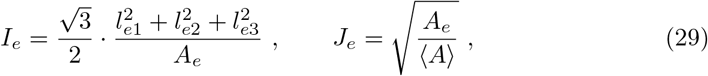

where *A_e_* is the area of the triangle *e* and *l*_*e*1_, *l*_*e*2_, *l*_*e*3_ its side lengths. *I_e_* and *J_e_* represent the invariants (trace and square root of the determinant) of the Cauchy-Green deformation tensor [59] assuming a reference equilateral triangle of size 〈*A*〉 ∑_*e*_ *A_e_/N_e_*, where the sum is taken over the *N_e_* triangles of a meshed surface. To specify the free energy *f*, we use the Neohookean energy *f*(*I_e_, J_e_*) = *μI_e_* + λ(*J_e_* – 1)^2^ [59]. Note that we define this energy on the mesh rather than on 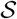. We want to minimise Eq. (28), but with the restriction that the cell shape 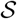 does not change, so that this operation corresponds to a surface reparametrisation without surface deformation. For this, we evolve the position of the vertices of the mesh according to velocities 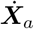. These velocities are obtained by introducing a continuous velocity field on the surface 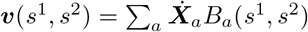, and by solving the following equations for the vertices velocities:

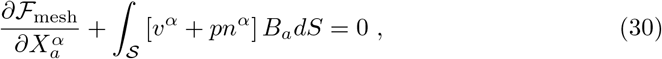

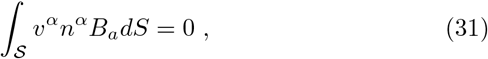

where *p*(*s*^1^, *s*^2^) = ∑_*a*_ *p_a_B_a_*(*s*^1^, *s*^2^) plays the role of a normal pressure, here a field enforcing the condition that the normal flow vanishes in a weak sense, Eq. (31). This leads to the relaxation of the free energy 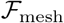, with vertices constrained to the shape of 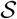, where the constraint is enforced weakly, i.e. in a finite element sense. In practice, this reparametrisation step is performed after a number of steps of the physical evolution of the cell surfaces, and we stop this mesh improvement dynamics when 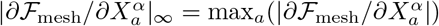 is smaller than a given tolerance.

## 3 Results

We now discuss applications of the interacting active surface framework. We first consider flows in a single spherical cell driven by gradient of cortical tension, a set-up which allows to compare simulation results to an analytical solution. We then examine the shape of the simplest multicellular aggregate, a doublet formed by two cells, when the two participating cells have equal or different tensions. Next, we consider an aggregate of cells assembled in a planar configuration, recapitulating the organisation of a small epithelial island. Finally, we introduce cell divisions in our framework and simulate the growth of a three-dimensional cell aggregate from a single cell. In the following we introduce a reference length scale 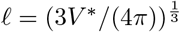 which corresponds to cell radius of a spherical cell, a reference surface tension 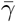, and a reference time scale 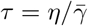, which corresponds to the characteristic time scale of cortical flows. We use these reference quantities for normalisation of other quantities.

### 3.1 Single cell: convergence with mesh size

We first consider flows driven by gradients of active tension in a single cell. This allows us to test the convergence of the numerical method for the dynamics of a single spherical cell, since we can compare the velocity field resulting from the method discretisation to an analytical solution using spherical harmonics. We consider a pattern of surface tension on a spherical surface of radius *ℓ*, given by

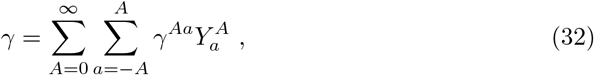

where *Y^Aa^* is the spherical harmonic of degree *A* and order *a*. The volume enclosed by the surface is assumed to be subjected to a uniform pressure difference *P*. The resulting velocity field can then be written as (S1 Appendix 7)

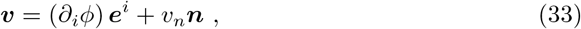

where the functions *ϕ* and *v_n_* can also be expanded in spherical harmonics, 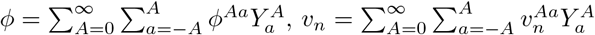, and the coefficients read:

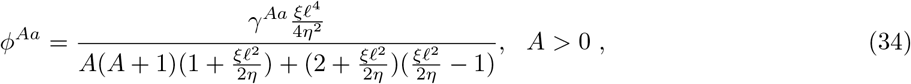

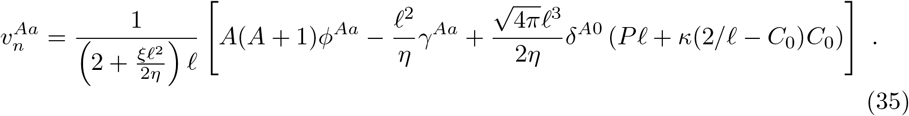

In Fig. 3, we consider flows resulting from a surface tension profile given by the coefficients 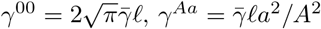 if 1 < *A* ≤ 4 and *γ^Aa^* = 0 for *A* > 4, and from the imposed inner pressure 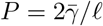 (Fig. 3A). We obtain the velocity field analytically (***v***^*^, Fig. 3B) and numerically (***v***, Fig. 3C), for different mesh sizes. We then compute the *L*_2_ norm of the error ***v*** – ***v***^*^, i.e. 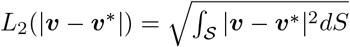, with respect to the *L*_2_ norm of ***v***^*^, 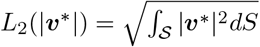. We find an excellent agreement between the exact and numerically obtained velocity field (Fig. 3B-D). The corresponding error scales with (*h/ℓ*)^2^, where *h* is the average mesh size, in line with the reported convergence rate of subdivision surfaces for other systems of partial differential equations [60] (Fig. 3D).

**Fig 3.**
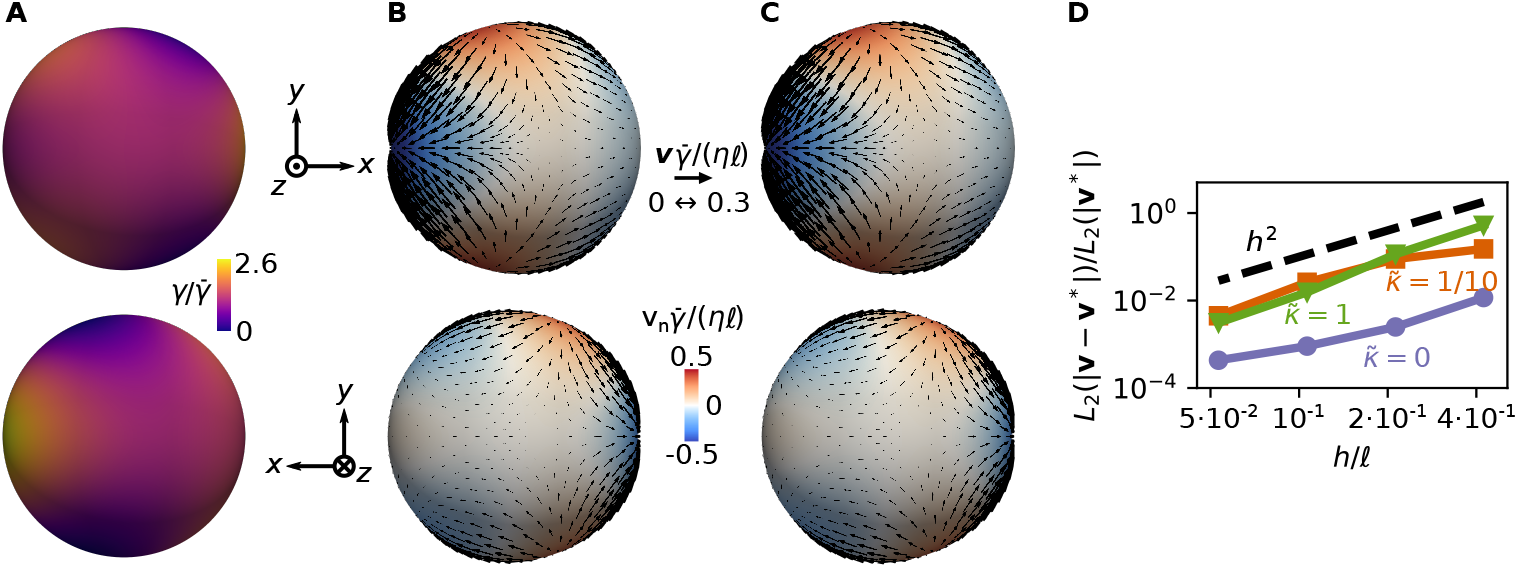
Single cell dynamics and convergence of the numerical framework. (A) An inhomogeneous pattern of active surface tension *γ* is imposed on a spherical surface. (B) Analytically computed velocity field generated by the active tension profile in A. The velocity field is decomposed into its normal (colormap) and tangential (arrows) components. (C) Numerical solution for the velocity profile generated by the active tension profile in A. (D) Discretisation error, evaluated here in terms of the *L*_2_ norm of the difference between the analytical and numerical solutions for the velocity, as a function of the average mesh size *h* and for 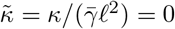 (blue), 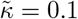 (orange) and 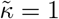 (green). Other parameters are *ξℓ*^2^/*η* = 4, *C*_0_*ℓ* = 1.

### 3.2 Shape and dynamics of an adhering cell doublet

We now discuss the equilibrium shape of an adhering cell doublet. In this and the following sections, the cell pressure difference *P_I_* is imposed as a Lagrange multiplier enforcing the condition *V_I_* = *V*_0_, with *V_I_* the cell volume and 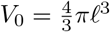 a reference volume, and we assume that *C*_0_ = 0. With these choices, 5 normalised, non-dimensional parameters have to be specified for each simulation: 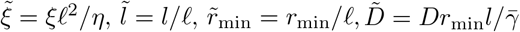, and 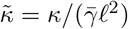.

In the following we set *ξ* = 10^-3^ so that the effect of friction is small. A cadherin bond has a typical length ~15-30 nm [61] and a typical actomyosin cortex thickness is 200nm [62], both much smaller than the typical radius of a cell, ~ from a few to tens of μm. Therefore, in the interaction of potential of cell surface we take 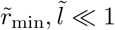. For simplicity, in the following we constrain 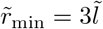. We typically choose values 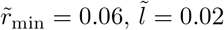 which for a cell radius of 5*μ*m, correspond to *r*_min_ = 300nm and *l* = 100nm.

We note that if the reference cell volume *V*_0_ is constant, the system can be viewed as a generalised gradient flow minimising the net free energy 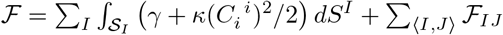, subjected to the constraint *V* = *V*_0_, where the first term represents an effective energy for the active tension *γ* and 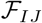 is defined in Eq. (16). We thus expect the system to eventually reach an equilibrium state with vanishing cortical flows.

We first analyse the behaviour of a doublet of identical cells. We initialise the simulation by putting two spherical cells close to each other, such that they are within the interaction range of the potential *φ*(*r*) (Eq. (26)) without touching. As expected, after an initial transient and contact growth, the doublet reaches an equilibrium shape (Fig. 4A-B). Increasing the relative adhesion strength 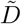 leads to an increasing adhesion patch and a lower cell pressure (Fig. 4C-D and S2 Fig. B). The value of the normalised distance 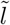 modulates the distance between the two cells (Fig. 4E). For 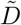 beyond a threshold which depends on 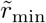 and 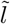, the adhesion patch develops a buckling instability. We found that this instability eventually leads to self-intersection of the computational mesh, as there is no energy contribution in our framework preventing such self-intersection (S2 Fig. A). Varying the bending modulus 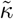, but maintaining relatively small values 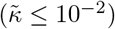, we find slightly smoother shapes at the boundary of the adhesion patch as well as a slight modulation of the threshold for buckling instability of the contact zone, with larger 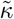 leading to higher stability (Figs. 4C,E).

**Fig 4.**
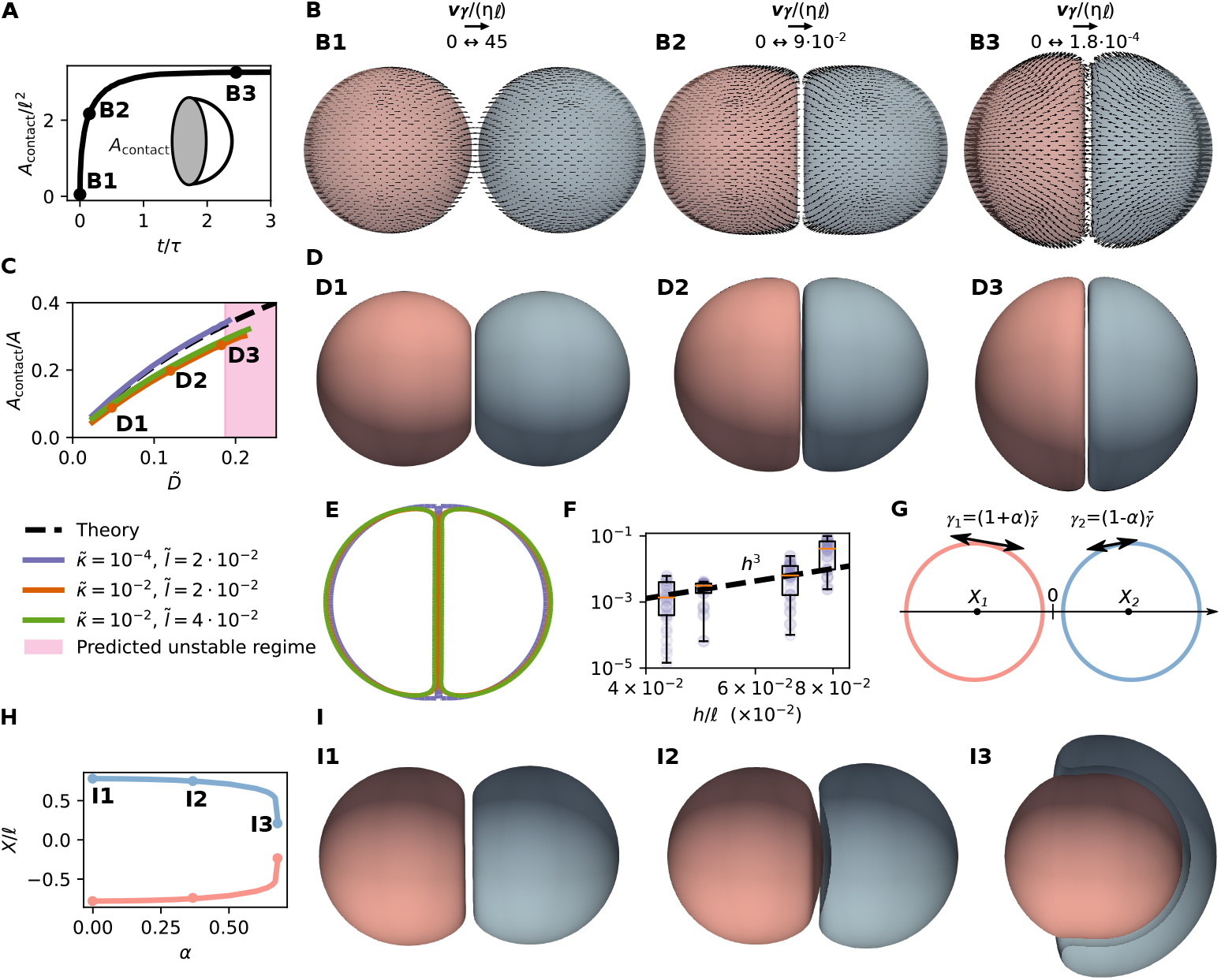
Dynamics and steady-state shape of a cell doublet. (A) Normalised area of contact as a function of time 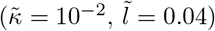. (B) Snapshots of deforming doublet at times indicated in (A), with the cell surface velocity field superimposed. (C) Coloured lines: ratio of contact surface area *A*_contact_ to cell surface area *A*, for simulations with different values of 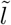 and 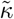. Black dotted line: theoretical approximation valid in the limit of 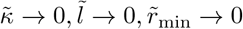. (D) Snapshots of doublet equilibrium shape for increasing adhesion strength; different parameters in (D1)-(D3) correspond to points labelled in C. (E) Comparison of a slice for the different simulations, with values of 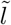 and 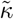 indicate in C, and 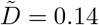. The value of 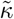 affects the shape smoothness of the edge of the adhesion patch. (F) Convergence of the method evaluated by computing the inner cell pressure *P* for different average mesh sizes *h*, and comparing the results with a simulation with *h/∓* ≈ 2 · 10^-2^ (finer). For each *h*, we compute a box plot using different values of 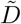 and fixed 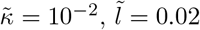. (G) Schematic of adhering doublet, with different active tensions *γ*^1^ and *γ*^2^ for each cell. (H) Position of the cell centre of mass *X*_1_ and *X*_2_, as a function of active tension asymmetry between the two adhering cells, *α* = (*γ*^1^ – *γ*^2^)/(*γ*^1^ + *γ*^2^), where *γ*^1^ and *γ*^2^ are the surface tensions of the two cells. Beyond *α* = 0.69, the cell with lowest tension completely engulfs the one with highest tension. (I) Snapshots of doublet equilibrium shape, clipped by a plane passing by the line joining the cell centres, for increasing difference of active surface tension; corresponding to points labelled in H. In H, I: 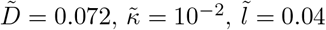.

To interpret these results, we turn to an approximate analysis of the shape of a doublet. Assuming a small bending rigidity 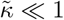, we approximate the cell doublet by two spherical caps of height *h_c_* and base of radius *r_c_*, forming an adhesion patch where the cells are separated by a distance *d* (Fig. 4E). The effective free energy of such a doublet configuration can then be written as

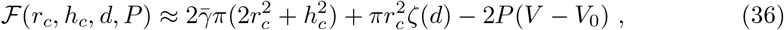

where 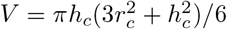 is the volume of one cell, *P* is the cell pressure, and *ζ*(*d*) an effective surface tension at the contact arising from cell-cell adhesion. For a sufficiently large patch compared to the interaction distances *r*_min_, and *l*, the effective surface tension can be approximated as (S1 Appendix 8):

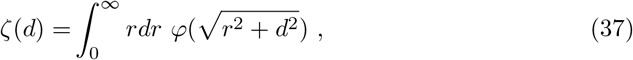

which can be evaluated numerically for a given potential *φ*. Minimising the effective free energy (36), one obtains equilibrium values 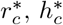, *d^*^, P^*^* (S1 Appendix 8), which depend on the effective surface tension of a cell at the contact:

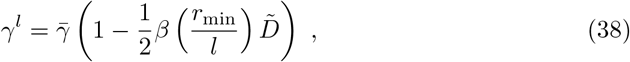

where the surface tension of the whole interface is 2*γ^l^*. Here, we have introduced a numerical function 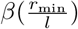, whose functional form depends on the potential *φ*(*r*). For the value of *r*_min_/*l* chosen here, *β* ≃ 10.7. When the net tension at the contact 2*γ^l^* becomes negative, for

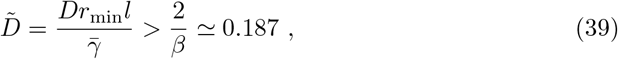

we expect the system to develop a buckling instability. Indeed, the corresponding threshold for buckling is well predicted by the simulation with smallest *l/ℓ* and bending modulus *κ* (Fig. 4C). Before the buckling instability, the ratio between the contact area and the cell surface area as well as the cell pressure *P^*^* are well predicted by the approximate analysis, which become more accurate as 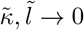 (Fig. 4C and S2 Fig. B).

To further check the numerical method, we analyse the convergence of the intracellular pressure *P* as a function of the mesh size *h*, by comparing the pressure value *P* obtained at different values of *h* and 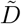 (for a fixed 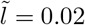) with a simulation with a fine mesh *h/ℓ* ≈ 2 · 10^-2^ (Fig. 4F). We observe that, on average, errors converge as ~ *h*^3^.

Finally, we examine an asymmetric doublet system where cells have different tensions *γ*^1^ and *γ*^2^ (Fig. 4G-I). The corresponding equilibrium state has been considered previously (Ref. [63] and references therein). Fixing a relatively low value of 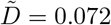, we change the ratio *α* = (*γ*^1^ – *γ*^2^)/(*γ*^1^ + *γ*^2^). As expected from previous studies, we observe that the cell with lowest tension progressively engulfs the cell with highest tension as the ratio *α* is increased. As a result, their centre of masses approach each other for increasing values of *α* (Fig. 4H-I). In our numerical simulations, beyond *α* ≃ 0.69, the cell with lowest tension self-intersects before completely engulfing the one with highest tension (S2 Fig. C).

### 3.3 Epithelial monolayer

We now discuss simulations of aggregates containing a larger number of cells. We start by considering a sheet of cells with free boundary conditions, intended to resemble the organisation of a simple epithelial island. To obtain an initial condition for such a sheet, spherical cells with homogeneous active tension 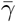 are positioned with their centres on a 6×6 hexagonal lattice with side *s* = 2*ℓ* + *r*_min_ + *l*. We then let the system relax to its equilibrium state, varying the adhesion parameter 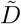, keeping its value below the instability threshold identified in the doublet analysis. As the adhesion parameter 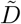 is increased, the cellular shapes progressively deviate from loosely adhering spheres to packed columnar cells (Fig. 5A-B). The reduced volume 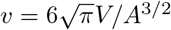 with *V* the cell volume and *A* its surface area, a measure of the deviation of the cell shape from a sphere with *v* = 1 being a sphere, is close to 1 for small 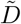 and then decreases (Fig. 5B).

**Fig 5.**
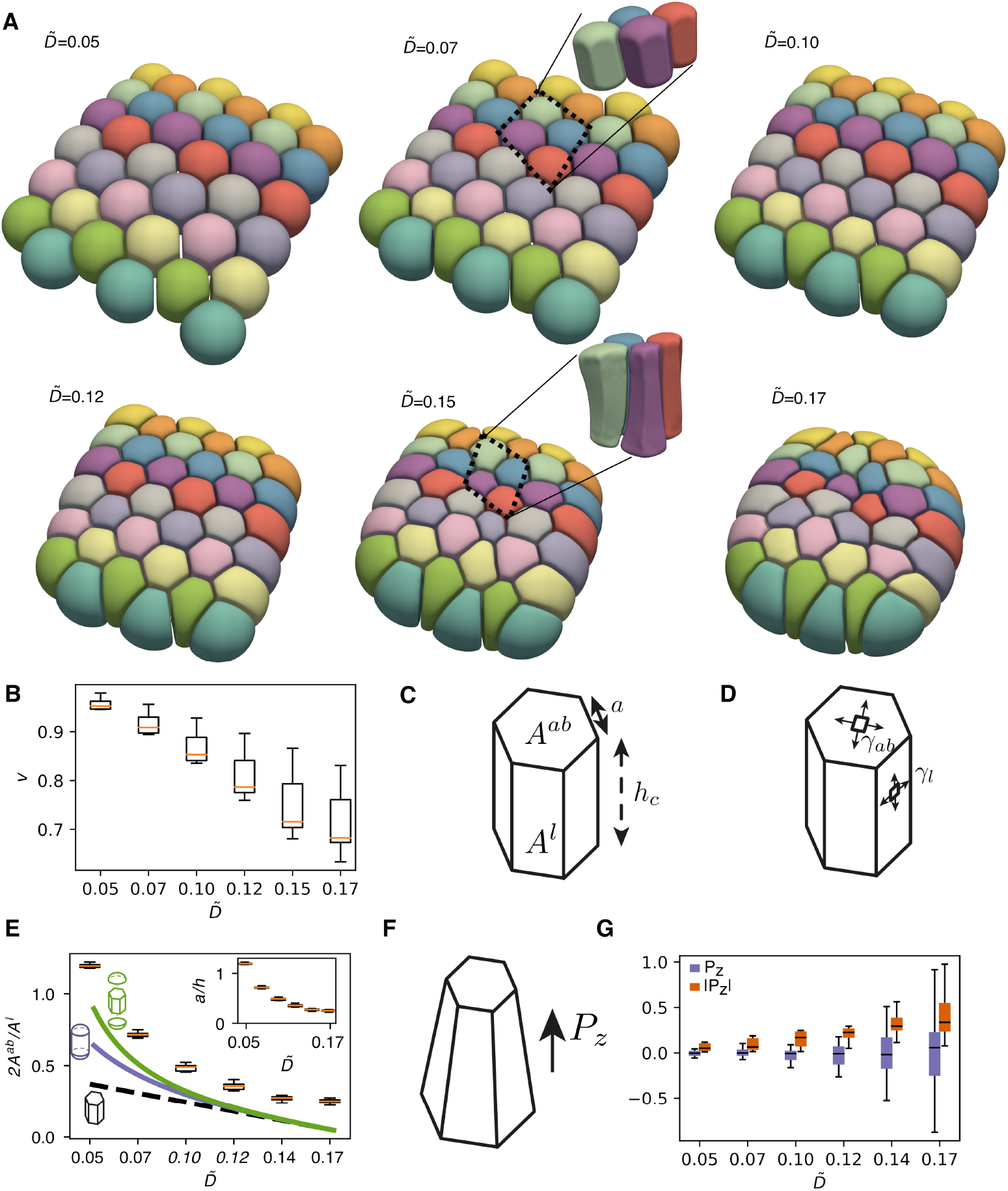
(A) Simulation results (equilibrium shapes) for a planar sheet for different values of the adhesion parameter 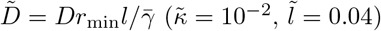. (B) Reduced volume *v* as a function of 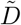. (C) Schematic for the measured apical and basal cell surface area *A^ab^*, lateral surface area *A^l^*, side length *a* and cell thickness *h_c_*. (D) Results are compared to a simple 3D vertex model with a lateral surface tension *γ_l_* and apical and basal surface tension *γ_ab_*. (E) Box plots: ratio of apico-basal to lateral surface area 2*A^ab^/A_l_* for the center cells, as a function of the adhesion parameter 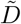. Dashed black, blue and green lines: prediction of simplified theories describing the cell shape as an hexagonal prism, a cylinder with two spherical caps, and the union of a hexagonal prism with two spherical caps. Inset: cellular aspect ratio *a/h_c_*, with *a* measuring the side of the hexagonal face and *h_c_* the thickness of the sheet, as a function of 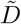. Here 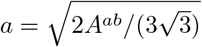 and *h* = *A^l^*/(6*a*). (F) Schematic for the polar vector ***P*** characterising the asymmetry of the cell shape. (G) Box plot for *P_z_* (blue) and |*P_z_*| (red). As adhesion increases, cells deform asymmetrically in the direction orthogonal to the planar sheet.

We compare the simulation results with a theoretical prediction from a 3D vertex model on a perfect hexagonal lattice, i.e. formed by uniform hexagonal prisms with height *h_c_* and side length *a* (Figs. 5C-D). We consider that cells are subjected to an apico-basal tension 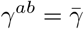, lateral tension 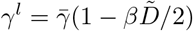, and an inner pressure *P* enforcing the cell volume to be equal to the reference volume *V*_0_ = 4*πℓ*^3^/3. We then write the corresponding effective free energy for a single cell in the tissue (S1 Appendix 9):

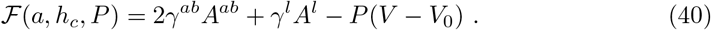

For a hexagonal prism, 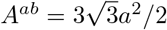 the apical and basal surface area, *A^l^* = 6*ah_c_* the lateral surface area, and the cell volume is 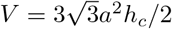. Minimising the effective free energy, we obtain the equilibrium ratio *A^ab^/A^l^* which can be compared to simulation results for the 16 inner cells (Fig. 5E). This area ratio is related to the cell aspect ratio, with smaller values corresponding to more columnar cells and larger values corresponding to more squamous cells. Although this simplified model captures the qualitative trend of the cell shape dependency on cell adhesion, it underestimates the simulated area ratio. We now consider alternative simplified descriptions where (i) each cell is considered as a cylinder connected to two spherical caps, or (ii) use an approximation where the surface area and volume of the cell is defined through the union of a hexagonal prism and two spherical caps (S1 Appendix 9). These choices better capture the actual simulated shapes (Fig. 5E), showing that taking the apical and basal curvatures play a significant role in the equilibrium cell shape. These refined models however still underestimate the area ratio *A^ab^/A^l^*. This can arise from the fact that these simplified descriptions do not take into account the surface bending modulus, the tissue-scale deformation due to the system finite size and free boundary conditions, and consider approximate cell shapes.

In addition, we noticed that for higher values of 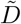, the cellular shapes appear more heterogeneous (Fig. 5A). This heterogeneity appears to be linked to an asymmetry in the cell apical and basal surface areas (see zoomed area comparing how lateral faces look for small 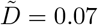 and for larger 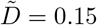 in Fig. 5A). To verify this, we introduce a polar order parameter for the shape of cell *I* (Fig. 5F):

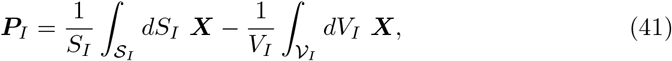

with 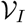 the volumetric domain enclosed by 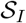, and *S_I_* and *V_I_* the surface area and volume of cell *I*. We calculate the order parameter for all cells in the simulation and consider its projection on the direction orthogonal to the plane containing the initial cell centers, *z*. We observe that, with increasing adhesion strength 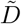, the average projected cell polarity does not clearly deviate from zero, 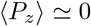, but the average of the absolute value of the projected cell polarity, 〈|*P_z_*|〉, strongly increases (Fig. 5G). This suggests that at high enough adhesion, cells adopt polarised apico-basal shapes orthogonal to the plane of the tissue, with no consistent overall shape polarisation orthogonal to the tissue (Fig. 5A). Possibly, such a spatial arrangement favours larger contact areas, which is beneficial at large adhesion.

### 3.4 Adding cell divisions: growth of an organoid

We now discuss simulations modelling the growth of a cell aggregate from a single cell, for which we introduce cell divisions in our framework. When a cell divides, the mother cell is replaced by two daughter cells as follows: a randomly oriented plane passing through the mother cell centre is selected, splitting the mother cell in two parts. The two daughter cells are then separated by this plane and simply fill the original shape of the mother, except for a small region that separates the daughter cells by a distance *d*\*, perpendicular to the division plane (Fig. 6A). To determine timepoints of cell division, each cell is assigned a cell cycle time *t_D_*, which we take equal for all cells. We assume that in between divisions, cell volume follows a linear growth law 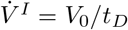, where *V*_0_ is the volume of the cell at its birth. This effectively leads to cells doubling their volume during their lifetime - we note however that a small volume loss occurs at division due to the initial separation of the daughter cells by a distance *d*\* (Fig. 6A). At each time point, cell volume is imposed through the Lagrange multiplier *P_I_*.

**Fig 6.**
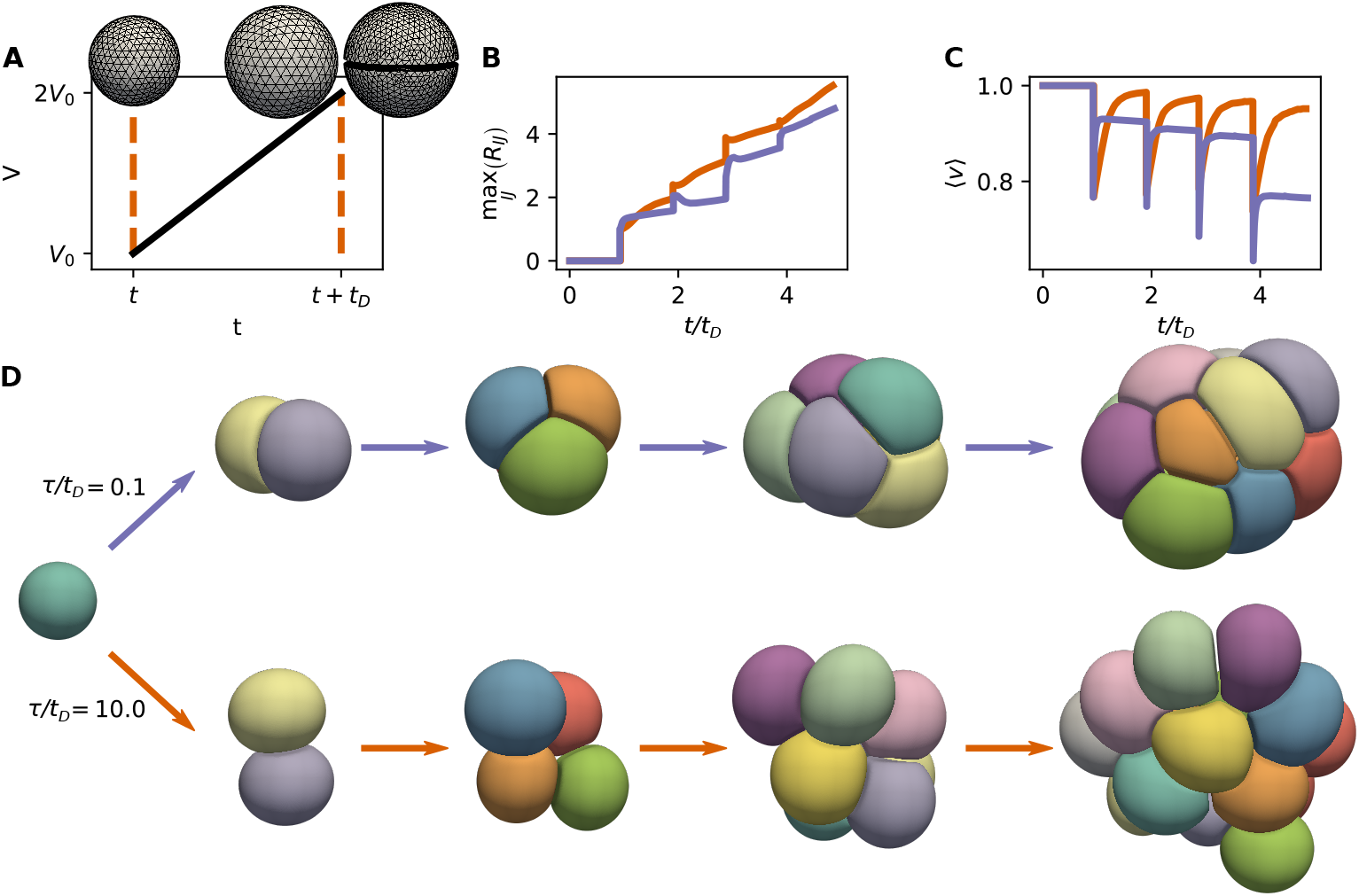
Growth of small cell aggregates, driven by synchronous cell divisions and cell volume increase 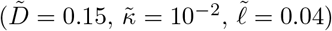. (A) Each cell doubles its volume between birth and division, over a cell cycle time *t_D_*. Cell division is introduced by splitting the mother cell with a plane passing through the cell centre, and generating two daughter cells separated by a distance *d*\* (*d*\*/2 from the division plane). (B-D) Simulation results for two values of the ratio *τ/t_D_*. (B): Largest centre-to-centre cell distance max〈*I,J*〉*R_IJ_*, as a function of time. Jumps correspond to cell division events. (C) Average reduced volume 〈*v*〉 as a function of time. (D) Snapshots of simulations of two growing aggregates.

We then simulate the growth of an aggregate starting from a single cell (Fig. 6B-D and S3-4 videos). The dynamics of the growing aggregate strongly depends on *τ/t_D_*, which measures the ratio between a characteristic time scale of cell shape relaxation, and the cell cycle time. For smaller values of *τ/t_D_*, the growing aggregate is more compact (Fig. 6B), cells have a smaller reduced volume and have therefore shapes further away from spheres (Fig. 6C). Here, the aggregate compactness is measured by calculating the maximum distance between cell centres (Fig. 6B). These observations indicate that the shape of a cell aggregate can strongly depend on a competition between its growth rate and internal mechanical relaxation times.

## 4 Discussion

The framework of interacting active surfaces introduced here is a novel method to study the mechanics of cell aggregates such as early developing embryos or organoids, and opens the door to their systematic modelling and simulation. We have demonstrated here that it can be used to study in detail the shape of adhering cell doublets, simple epithelia, as well as growing cellular aggregates. Our method is well-suited to capture the mechanics of tissues and organoids connecting it to cell level processes such as cortical flows, cortical tension and cellular adhesion in the organisation of a cellular aggregate.

In this study we have restricted ourselves to relatively simple constitutive equations for the tension and bending moment tensors (Eqs. (3)), and we have considered situations with a uniform and constant surface tension within each cell. Our method is based on using the virtual work principle (Eq. (2)), a very general statement of force and torque balance for a surface, to obtain a set of algebraic equations for the cell surface described with finite elements. As such it is versatile and we expect that more complex constitutive equations, corresponding to more detailed physical descriptions of the cell surface, can be easily introduced in our description. We now discuss some of these possible extensions of our framework.

First, we have not included here apico-basal polarity, an axis of cell organisation which results from a spatially segregated protein distribution and inhomogeneous cytoskeletal structures [64]. To take this into account, one could introduce a polarity field in each cell and consider an active tension γ on the cell surface whose value at each point depends on the polarity field orientation. This could be used to introduce, for instance, differences in apical, basal and lateral surface tension which are taken into account in 3D vertex models [19].

It would be natural to introduce a concentration field on the cell surface, describing a regulator of the cortical tension, such as myosin concentration. At the level of a single surface, such coupling between cortical flows and its regulator can give rise to pattern formation, spontaneous symmetry breaking and shape oscillation [37, 65, 66]. The dynamics of the concentration per unit area, *c*, of such a regulator can be obtained from the balance equation on the surface:

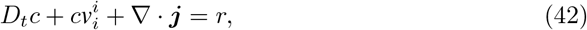

where *D_t_c* is the material derivative of *c* (*∂_t_c* in a Lagrangian description), ***j*** is the flux of c relative to the centre of mass, and *r* is a reaction rate. A natural choice for the flux would be **j** = –*D***∇***c* to represent diffusion according to Fick’s law. A natural choice for the reaction rate would be *r* = *k*_on_ – *k*_off_*c* for turnover dynamics, with target concentration *c*_0_ = *k*_on_/*k*_off_ and typical turnover time 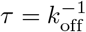. The discretisation of such fields can be easily introduced in our framework, following the methods detailed in [38,67].

Importantly, the model introduced here is based on an interaction potential between cells, which can be motivated microscopically from a description of cell-cell linkers that equilibrate quickly to their Boltzmann distribution, with free linkers on the surface in contact with a reservoir imposing a constant concentration. From a computational perspective, the exact integration of the interaction potential requires the computation of double integrals, which have a large computational cost. Alternatively, one could approximate the double integrals further in the limit 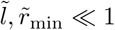 by considering the interaction of each point on surface 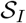 only with its closest point projection on 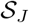, following classical numerical approaches for the adhesion between interfaces (Ref. [68] and references therein).

On the other hand, the interplay of adhesive cell-cell linkers such as E-cadherin with cortical dynamics also plays an important role in orchestrating cell adhesion [69]. Unlike the specific adhesion of solid interfaces, cell-cell adhesion dynamics involves a complex interplay between the diffusion, advection and binding dynamics of linkers [70]. Notably, E-cadherin junctions have been shown to be mechanosensitive [71] and to act to regulate the actomyosin levels at junctions [72]. To take these effects into account, an explicit description of E-cadherin concentration on the cell surface might be required. Thus, an extension of our model could introduce explicitly two-point density fields *c_IJ_* (***X***_1_, ***X***_2_) representing the concentration of bound linkers between cells *I* and *J*, as well as a concentration field of free linkers on each cell *c_I_*. Alternatively, one could introduce cell-cell adhesion by considering a finite number of explicitely described individual linkers [73].

Our model does not account for the friction generated by relative surface flows between cells that adhere to each other, which is likely to play an important role during cell rearrangements. The effective friction stemming from an ensemble of transiently binding and unbinding linkers can be modelled effectively with a friction coefficient motivated by microscopic models such as a Lacker-Peskin model [74], which lead to predictions of force-velocity relations which depend on whether linkers are force-sensitive, e.g. slip or catch bonds [75, 76]. One could include these terms systematically in our finite element discretisation following the ideas in [77, 78].

In its current version and with these additions, we hope that the interacting active surface framework will be a useful tool to investigate the mechanics and self-organisation of cellular aggregates.

## 5 Author contributions

Conceptualisation, Methodology, Writing -Original draft preparation, Writing -Review and editing: ATS, GS. Software: ATS. Validation: MKW. Funding acquisition, Supervision: GS.

## 6 Acknowledgements

ATS, MKW and GS acknowledge support from the Francis Crick Institute, which receives its core funding from Cancer Research UK (FC001317), the UK Medical Research Council (FC001317), and the Wellcome Trust (FC001317). GS and ATS acknowledge support from a grant from the Engineering and Physical Sciences Research Council (EP/T003103/1). We thank Quentin Vagne and Guillermo Vilanova for comments on the manuscript.

