## Supplementary material for "Interacting active surfaces: a model for three-dimensional cell aggregates": S1 Appendix

March 21, 2022

#### 1 Notations of differential geometry

Here we summarise notations of differential geometry used in this study, which follow Refs. [1] and [2]. The surface $\mathcal{S}$ is described by the parametrisation $\boldsymbol{X}(s^1, s^2)$, with $\boldsymbol{X}$ a 3D vector and $s^1$, $s^2$ two surface coordinates. The cartesian basis of the 3D space in which the surface is embedded is denoted $\tilde{\boldsymbol{e}}_\alpha$, with $\alpha = x, y, z$. In the following we use greek indices to label 3D coordinates, and latin indices to label coordinates on the surface. The tangent vectors $\boldsymbol{e}_i$, normal vector $\boldsymbol{n}$, metric and curvature tensors $g_{ij}$ and $C_{ij}$ and Christoffel symbols $\Gamma^k_{ij}$ are respectively defined by:

$$\boldsymbol{e}_i = \partial_i \boldsymbol{X} \ , \tag{1}$$

$$\boldsymbol{n} = \frac{\boldsymbol{e}_1 \times \boldsymbol{e}_2}{|\boldsymbol{e}_1 \times \boldsymbol{e}_2|} \ , \tag{2}$$

$$g_{ij} = \boldsymbol{e}_i \times \boldsymbol{e}_j \ , \tag{3}$$

$$\partial_i \boldsymbol{e}_j = -C_{ij}\boldsymbol{n} + \Gamma^k_{ij}\boldsymbol{e}^k \ . \tag{4}$$

The determinant of the metric tensor is denoted $g = \det g_{ij}$, and the surface element is $dS = \sqrt{g} ds^1 ds^2$. The vectors $\boldsymbol{e}^1$, $\boldsymbol{e}^2$ of the dual basis are defined by:

$$\boldsymbol{e}_i \cdot \boldsymbol{e}^j = \delta_i^j \ . \tag{5}$$

The inverse of the metric tensor $g_{ij}$ is denoted $g^{ij}$, such that $g_{ik}g^{kj} = \delta_i{}^j$. We also introduce the Levi-Civita tensor on a curved surface:

$$\epsilon_{ij} = (\boldsymbol{e}_i \times \boldsymbol{e}_j) \cdot \boldsymbol{n} \ , \tag{6}$$

which verifies $\epsilon_{ij}\epsilon^j{}_k = -g_{ik}$.

A 3D vector $\boldsymbol{a} = a^\alpha \tilde{\boldsymbol{e}}_\alpha$ has tangential and normal components to the surface, according to:

$$\boldsymbol{a} = a^i \boldsymbol{e}_i + a_n \boldsymbol{n} = a_i \boldsymbol{e}^i + a_n \boldsymbol{n} \ . \tag{7}$$

The metric tensor $g_{ij}$ and its inverse $g^{ij}$ can be used to raise and lower indices of vector and tensor components of the surface; for instance $a^i = g^{ij} a_j$ and $a_i = g_{ij} a^j$.

We also introduce covariant derivatives of a tangent vector $\boldsymbol{a}$ and tangent second-rank tensor $\boldsymbol{b}$ on the surface as follows:

$$\nabla_i a^j = \partial_i a^j + \Gamma^j_{ik} a^k \ , \tag{8}$$

$$\nabla_i t^{jk} = \partial_i t^{jk} + \Gamma^j_{il} t^{lk} + \Gamma^k_{il} t^{jl} \ . \tag{9}$$

The covariant derivatives of the metric and Levi-Civita tensor vanish, $\nabla_i g_{jk} = 0$ and $\nabla_i \epsilon_{jk} = 0$. In the following we also use the notation $\partial^i = g^{ij}\partial_j$ and $\nabla^i = g^{ij}\nabla_j$.

#### 2 Variations of the metric, curvature and Christoffel symbols

Given a variation of the surface parametrisation $\delta\boldsymbol{X}$, we can compute the variation of the different geometrical quantities of the surface. For the tangent vectors, we have

$$\delta \boldsymbol{e}_i = \delta \partial_i \delta \boldsymbol{X} = \partial_i \delta \boldsymbol{X} \ . \tag{10}$$

The variation of the metric tensor can then be computed as:

$$\delta g_{ij} = \delta(\boldsymbol{e}_i \cdot \boldsymbol{e}_j) = \delta\boldsymbol{e}_i \cdot \boldsymbol{e}_j + \boldsymbol{e}_i \cdot \delta\boldsymbol{e}_j = \partial_i \delta\boldsymbol{X} \cdot \boldsymbol{e}_j + \boldsymbol{e}_i \cdot \partial_j \delta\boldsymbol{X} \ . \tag{11}$$

For the inverse of the metric, we can use that $g^{ij} g_{jk} = \delta^i_k$, and then

$$\delta(g^{ij} g_{jk}) = 0 \Longrightarrow \delta g^{ij} = -g^{ik} g^{jl} \delta g_{kl} \ . \tag{12}$$

The variation of the metric determinant $g = \det(g_{ij})$ is given by Jacobi's formula:

$$\delta g = g g^{ij} \delta g_{ij} \ . \tag{13}$$

As a result, we have the useful relation for the infinitesimal variation of the square root of the metric determinant:

$$\delta\left(\sqrt{g}\right) = \frac{1}{2} g^{ij} \delta g_{ij} \sqrt{g} \ . \tag{14}$$

Eq. (12) can be used then to compute variations of the basis vectors of the dual basis

$$\delta\boldsymbol{e}^i = \delta\left(g^{ij} \boldsymbol{e}_j\right) = \delta g^{ij} \boldsymbol{e}_j + g^{ij} \partial_j \delta\boldsymbol{X} \ . \tag{15}$$

To compute variations of the normal, we note that $\boldsymbol{n} \cdot \boldsymbol{e}_i = 0$ and $\boldsymbol{n} \cdot \boldsymbol{n} = 1$ so $\delta\boldsymbol{n} \cdot \boldsymbol{e}_i = -\delta\boldsymbol{e}_i \cdot \boldsymbol{n}$, and $\delta\boldsymbol{n} \cdot \boldsymbol{n} = 0$, leading to:

$$\delta\boldsymbol{n} = -\left(\delta\boldsymbol{e}_i \cdot \boldsymbol{n}\right) \boldsymbol{e}^i = -\left(\partial_i \delta\boldsymbol{X} \cdot \boldsymbol{n}\right) \boldsymbol{e}^i \ . \tag{16}$$

The variations of the curvature tensor are then,

$$\delta C_{ij} = -\delta\left(\boldsymbol{n} \cdot \partial_i \boldsymbol{e}_j\right) = -\delta\boldsymbol{n} \cdot \partial_i \boldsymbol{e}_j - \boldsymbol{n} \cdot \partial_i \partial_j \delta\boldsymbol{X} \ . \tag{17}$$

Finally, variations of the Christoffel symbols have the form

$$\delta\Gamma^k_{ij} = \delta\left(\boldsymbol{e}^k \cdot \partial_i \boldsymbol{e}_j\right) = \delta\boldsymbol{e}^k \cdot \partial_i \boldsymbol{e}_j + \boldsymbol{e}^k \cdot \partial_i \partial_j \delta\boldsymbol{X} \ . \tag{18}$$

Expressions (11), (17), and (18) can be rewritten in terms of infinitesimal tangential displacement $\delta X^i$ and normal displacement $\delta X_n$ by substituting $\delta\boldsymbol{X} = \delta X^i \boldsymbol{e}_i + \delta X_n \boldsymbol{n}$. This leads to the expressions [1]

$$\delta g_{ij} = \nabla_i \delta X_j + \nabla_j \delta X_i + 2C_{ij} \delta X_n \ , \tag{19}$$

$$\delta C_{ij} = \delta X^l \nabla_j C_{il} - \nabla_j \nabla_i \delta X_n + C^l_i C_{jl} \delta X_n + C_{lj} \nabla_i \delta X^l + C_{il} \nabla_j \delta X^l \ , \tag{20}$$

$$\begin{aligned} \delta\Gamma^k_{ij} &= (\nabla_j C_i{}^k) \delta X_n + C_j{}^k \partial_i \delta X_n + C_i{}^k \partial_j \delta X_n - C_{ij} \partial^k \delta X_n \\ &\quad + \nabla_j \nabla_i \delta X^k + \left(C_l{}^k C_{ij} - C_{il} C_j{}^k\right) \delta X^l \ , \end{aligned} \tag{21}$$

which can be used to obtain the virtual work expression Eq. (34).

When the surface changes according to a flow field $\boldsymbol{v}$, and coordinates follow the flow in a Lagrangian way, the surface shape change can be written $\delta X = \boldsymbol{v} dt$. Therefore, using Eq.(19), the rate of change of the components of the metric tensor is:

$$\frac{dg_{ij}}{dt} = \nabla_i v_j + \nabla_j v_i + 2C_{ij} v_n = 2v_{ij} \ , \tag{22}$$

where $v_{ij}$ is the strain-rate tensor, defined in Eq. (4) in the main text. Eq. (22) is the definition for the components of the Lie derivative of $\boldsymbol{g}$ along the flow $\boldsymbol{v}$ [3].

### 3 Balance of linear and angular momentum and the principle of virtual work

Here we show that the condition of vanishing infinitesimal virtual work, Eq. (2) in the main text, follows from the force balance and torque balance equations on the surface $\mathcal{S}$, at low Reynolds number where inertial terms can be neglected. We consider a more general case than in the main text, where the surface subjected to an external force density $\boldsymbol{f}$ and an external torque density $\boldsymbol{\Gamma}$. Internal forces and torques are described by the tension tensor $\boldsymbol{t}^i = t^{ij} \boldsymbol{e}_j + t^i_n \boldsymbol{n}$ and the moment tensor $\boldsymbol{m}^i = m^{ij} \boldsymbol{e}_j + m^i_n \boldsymbol{n}$, such that:

$$\boldsymbol{t}^i dl \boldsymbol{\nu}_i \ , \qquad \boldsymbol{m}^i dl \boldsymbol{\nu}_i \ , \tag{23}$$

are respectively the force and torque acting on a cut on the surface with length $dl$, with $\boldsymbol{\nu}$ the unit vector tangent to the surface and normal to the cut. The statement of conservation of linear momentum (or force balance) for a thin sheet, at low Reynolds number where inertial terms are neglected, is [1]

$$\nabla_i t^{ij} + C_i{}^j t^i_n + f^j = 0 \ , \tag{24}$$

$$\nabla_i t^i_n - C_{ij} t^{ij} + f_n = 0 \ , \tag{25}$$

where the first equation stems from balance of forces tangent to $\mathcal{S}$ and the second from balance of forces normal to $\mathcal{S}$. On the other hand, balance of angular momentum (or torque balance) tangent and normal to $\mathcal{S}$ reads

$$\nabla_i m^{ij} + C_i{}^j m^i_n - \epsilon_i{}^j t^i_n + \Gamma^j = 0 \ , \tag{26}$$

$$\nabla_i m^i_n - C_{ij} m^{ij} + \epsilon_{ij} t^{ij} + \Gamma_n = 0 \ . \tag{27}$$

In the following, for convenience we introduce the symmetric and antisymmetric part of the tension tensor $t^{ij}$:

$$t^{ij} = t^{ij}_S + t^{ij}_A \ , \tag{28}$$

where $t^{ij}_S = t^{ji}_S$ and $t^{ij}_A = -t^{ji}_A$. Eqs. (26) and (27) can be rewritten as:

$$t^i_n = \epsilon^i{}_k \nabla_j m^{jk} + \epsilon^i{}_k C_j{}^k m^j_n + \epsilon^i{}_k \Gamma^k \ , \tag{29}$$

$$t^{ij}_A = \frac{1}{2}\left[-\nabla_k m^k_n + C_{kl} m^{kl} - \Gamma_n\right] \epsilon^{ij} \ . \tag{30}$$

We now multiply Eq. (24) by an infinitesimal tangential displacement field $-\delta X_j$ and integrate the result over $\mathcal{S}$, to obtain:

$$\begin{aligned} 0 &= \int_{\mathcal{S}} dS(-\delta X_j)\left(\nabla_i t^{ij} + C_i{}^j t^i_n + f^j\right) \\ &= \int_{\mathcal{S}} dS\Big[\left(\nabla_i \delta X_j\right) t^{ij} + \epsilon^i{}_k \nabla_l\left(\delta X_j C_i{}^j\right) m^{lk} - m^l_n \epsilon^i{}_k C_l{}^k C_i{}^j \delta X_j \\ &\qquad - \epsilon^i{}_k C_i{}^j \Gamma^k \delta X_j - f^j \delta X_j\Big] \\ &= \int_{\mathcal{S}} dS\left\{(\nabla_i \delta X_j) t^{ij}_S + \left[\epsilon^i{}_k \nabla_l\left(\delta X_j C_i{}^j\right) + \frac{1}{2} C_{lk}\epsilon^{ij}\nabla_i \delta X_j\right] m^{lk}\right. \\ &\qquad + \left[-\epsilon^k{}_l C_i{}^l C_k{}^j \delta X_j + \frac{1}{2}\epsilon^{jk}\nabla_i \nabla_j \delta X_k\right] m^i_n \\ &\qquad \left. -\delta X_j f^j - \epsilon^i{}_k C_i{}^j \Gamma^k \delta X_j - \frac{1}{2}\left(\nabla_i \delta X_j\right)\Gamma_n\right\} , \end{aligned} \tag{31}$$

where we have used Eqs. (29) and (30) and integrated by parts terms taking into account that $\mathcal{S}$ is a closed surface. Multiplying Eq. (25) by an infinitesimal normal displacement field $-\delta X_n$ and again integrating over $\mathcal{S}$, we obtain:

$$\begin{aligned} 0 &= \int_{\mathcal{S}} dS(-\delta X_n)\left(\nabla_i t^i_n - C_{ij} t^{ij}_S + f_n\right) \\ &= \int_{\mathcal{S}} dS\Big[-\epsilon^i{}_k(\nabla_j(\partial_i \delta X_n)) m^{jk} - (\partial_j \delta X_n)\epsilon_k^j C_i{}^k m^i_n \\ &\qquad + (\partial_i \delta X_n)\epsilon^i{}_k \Gamma^k + \delta X_n C_{ij} t^{ij}_S - \delta X_n f_n\Big] \ . \end{aligned} \tag{32}$$

Adding Eqs. (31) and (32), we get

$$\begin{aligned} 0 = \int_{\mathcal{S}} dS &\left\{\frac{1}{2}\left(\nabla_i \delta X_j + \nabla_j \delta X_i + 2\delta X_n C_{ij}\right) t^{ij}_S\right. \\ &+ \left[\epsilon^i{}_k \nabla_l\left(\delta X_j C_i{}^j\right) + \frac{1}{2} C_{lk}\epsilon^{ij}\nabla_i \delta X_j - \epsilon^i{}_k \nabla_l(\partial_i \delta X_n)\right] m^{lk} \\ &+ \left[\frac{1}{2}\epsilon^{jk}\nabla_i \nabla_j \delta X_k + \epsilon^j{}_k C_i{}^k \partial_j \delta X_n + \epsilon_j{}^k C_i{}^j C_k{}^l \delta X_l\right] m^i_n \\ &\left. -\delta X_i f^i - \delta X_n f_n + \left[\partial_i \delta X_n - \delta X_j C_i{}^j\right]\epsilon^i{}_k \Gamma^k - \frac{1}{2}(\epsilon^{ij}\nabla_i \delta X_j)\Gamma_n\right\} . \end{aligned} \tag{33}$$

Finally, using the analytical expressions for the variations of the metric, curvature, and Christoffel symbols $\delta g_{ij}$, $\delta C_{ij}$, and $\delta \Gamma^k_{ij}$ in terms of tangential and normal variations (Eqs. (19)-(21)), we get

$$\delta W = \int_{\mathcal{S}} dS\left\{\frac{1}{2}\hat{t}^{ij}\delta g_{ij} + \bar{m}^{ij}\delta C_{ij} + \frac{1}{2} m^i_n \delta\Gamma^k_{ij}\epsilon^j{}_k - f^\alpha \delta X^\alpha - \frac{1}{2}\Gamma^\alpha\left(\nabla \times \delta \boldsymbol{X}\right)^\alpha\right\} = 0 \ , \tag{34}$$

where we have defined $\nabla \times \delta \boldsymbol{X} = 2\epsilon^{ij}(\partial_j \delta X_n - C_{jk}\delta X^k)\boldsymbol{e}_i + (\nabla_i \delta X_j)\epsilon^{ij}\boldsymbol{n}$ (see Refs. [1, 2]), $\bar{m}^{ij} = -m^{ik}\epsilon_k^{\ j}$, and the tension tensor

$$\hat{t}^{ij} = t_S^{ij} - \frac{1}{2}\left(\bar{m}^{ki}C_k^{\ j} + \bar{m}^{kj}C_k^{\ i}\right) , \tag{35}$$

which accounts for the work generated by the in-plane tension tensor $t^{ij}$ as well as the work generated by the bending moments $m^{ij}$ upon an in-plane deformation characterised by $\delta g_{ij}$. The expression for the differential virtual work, Eq. (2) in the main text, corresponds to Eq. (34) with vanishing external torque density ($\boldsymbol{\Gamma} = 0$) and vanishing normal part of the moment tensor ($m_n^i = 0$).

### 4 Equilibrium tension and bending moments in the Helfrich model

Here we derive the tension $\hat{t}_e^{ij}$ and bending moment $\bar{m}_e^{ij}$ arising from a Helfrich energy of the form

$$\mathcal{F}_{\text{Helfrich}} = \int_{\mathcal{S}} dS \frac{\kappa}{2}(C_i^{\ i} - C_0)^2 . \tag{36}$$

To identify $\hat{t}_e^{ij}$ and $\bar{m}_e^{ij}$, we take variations of Eq. (36),

$$\delta\mathcal{F}_{\text{Helfrich}} = \int_{\mathcal{S}} dS \left[\kappa(C_i^{\ i} - C_0)\delta\left(C_i^{\ i}\right) dS + \frac{\kappa}{2}\left(C_i^{\ i} - C_0\right)^2 \frac{\delta\sqrt{g}}{\sqrt{g}}\right] , \tag{37}$$

where we have used that $dS = \sqrt{g}ds^1 ds^2$ with $g$ the determinant of the metric. The differential of the trace of the curvature tensor is:

$$\delta\left(C_i^{\ i}\right) = \delta(g^{ij}C_{ij}) = C_{ij}\delta g^{ij} + g^{ij}\delta C_{ij} = -C^{ij}\delta g_{ij} + g^{ij}\delta C_{ij} , \tag{38}$$

where we have used that $\delta g^{ij} = -g^{ik}g^{jl}\delta g_{kl}$. Noting that the variation of the square root of the metric determinant is given by Eq. (14), we can then write:

$$\delta\mathcal{F}_{\text{Helfrich}} = \int_{\mathcal{S}} dS \left\{\frac{\kappa}{2}(C_k^{\ k} - C_0)\left[\frac{1}{2}(C_l^{\ l} - C_0)g^{ij} - 2C^{ij}\right]\delta g_{ij} + \kappa(C_k^{\ k} - C_0)g^{ij}\delta C_{ij}\right\}, \tag{39}$$

which, by comparing with Eq. (34) and using that $\delta W = \delta\mathcal{F}$ at equilibrium, allows us to identify

$$\hat{t}_e^{ij} = \kappa(C_k^{\ k} - C_0)\left[\frac{1}{2}(C_k^{\ k} - C_0)g^{ij} - 2C^{ij}\right], \qquad \bar{m}_e^{ij} = \kappa(C_k^{\ k} - C_0)g^{ij} , \tag{40}$$

which contribute to the total tension and bending moments in the constitutive equations, Eq. (3) in the main text.

### 5 Microscopic motivation for the cell-cell adhesion potential

Here we discuss a microscopic motivation for the characterisation of cell-cell interactions with a potential. Following Eq. 22 in the main text, we consider the free energy of an ensemble of linkers, described by a two-point concentration field $c_{IJ}(\boldsymbol{X}_I, \boldsymbol{X}_J)$ and concentration $c_I$ denotes the concentration of unbound, free linkers in cell $I$:

$$\begin{aligned}\mathcal{F}^{\text{micro}} =& \sum_I \int_{\mathcal{S}_I} dS_I k_{\text{B}}T c_I \left(\log\frac{c_I}{c_0} - 1\right) \\ &+ \sum_{\langle I,J\rangle} \int_{\mathcal{S}_I} dS_I \int_{\mathcal{S}_J} dS_J c_{IJ}\left[k_{\text{B}}T\left(\log\frac{c_{IJ}}{c_0^2} - 1\right) + \phi\left(|\boldsymbol{X}_I - \boldsymbol{X}_J|\right)\right] ,\end{aligned} \tag{41}$$

where parameter definitions are discussed in the main text.

Variation of the free energy (41) with respect to the concentrations gives:

$$\begin{aligned}\delta\mathcal{F}^{\text{micro}} =& \sum_I \int_{\mathcal{S}_I} dS_I k_{\text{B}}T \delta c_I \log\frac{c_I}{c_0} \\ &+ \sum_{\langle I,J\rangle} \int_{\mathcal{S}_I} dS_I \int_{\mathcal{S}_J} dS_J \delta c_{IJ}\left[k_{\text{B}}T\log\frac{c_{IJ}}{c_0^2} + \phi\left(|\boldsymbol{X}_I - \boldsymbol{X}_J|\right)\right] ,\end{aligned} \tag{42}$$

To obtain chemical equilibrium, we note that concentrations can change due to the reaction of binding of two linkers on surfaces $I$, $J$ with reaction coordinate $r_{IJ} = r_{JI}$, and can change due to exchange of free linkers with the bulk, with reaction coordinates $r_{0I}$:

$$\delta c_{IJ} = \delta r_{IJ} \ , \tag{43}$$

$$\delta c_I = -\sum_{J \neq I} \delta r_{IJ} + \delta r_{0I} \ , \tag{44}$$

such that minimisation with respect to $\delta r_{IJ}$ gives:

$$c_{IJ} = c_I c_J \exp\left[-\frac{\phi\left(|\boldsymbol{X}_I - \boldsymbol{X}_J|\right)}{k_{\mathrm{B}}T}\right] \ . \tag{45}$$

and minimisation with respect to $\delta r_{0I}$ gives $c_I = c_0$. One then obtains:

$$c_{IJ} = c_0^2 \exp\left[-\frac{\phi\left(|\boldsymbol{X}_I - \boldsymbol{X}_J|\right)}{k_{\mathrm{B}}T}\right] \ , \tag{46}$$

such that $\exp\left[-\frac{\phi(|\boldsymbol{X}_I - \boldsymbol{X}_J|)}{k_{\mathrm{B}}T}\right]$ is the binding affinity of linkers for two element of surface separated by the distance $|\boldsymbol{X}_I - \boldsymbol{X}_J|$.

Taking into account that a surface deformation dilutes locally the density of linkers as $\delta c_I = -\frac{1}{2} c_I g_I^{ij} \delta g_{I,ij}$ and $\delta c_{IJ} = -c_{IJ}\left(\frac{1}{2} g_I^{ij} \delta g_{I,ij} + \frac{1}{2} g_J^{ij} \delta g_{J,ij}\right)$, the variation of the free energy (41) with respect to a shape change leads to

$$\delta \mathcal{F}^{\mathrm{micro}} = \sum_I \int_{\mathcal{S}_I} dS_I \left(-\frac{1}{2} k_{\mathrm{B}} T g_I^{ij} c_I\right) \delta g_{I,ij} + \sum_{\langle I, \rangle J} \delta \mathcal{F}_{IJ} \tag{47}$$

with

$$\delta \mathcal{F}_{IJ} = k_{\mathrm{B}} T \int_{\mathcal{S}_I} dS_I \int_{\mathcal{S}_J} dS_J c_{IJ} \Big[ -\frac{1}{2}\left(g_I^{ij} \delta g_{I,ij} + g_J^{ij} \delta g_{J,ij}\right) + \phi'\left(|\boldsymbol{X}_I - \boldsymbol{X}_J|\right) \frac{X_I^\alpha - X_J^\alpha}{|\boldsymbol{X}_I - \boldsymbol{X}_J|} (\delta X_I^\alpha - \delta X_J^\alpha) \Big] \ . \tag{48}$$

We now assume that linkers can bind and unbind the surfaces quickly so that $c_I$ and $c_{IJ}$ are equal to their equilibrium values. The contribution to the first sum in Eq. (47) is then equivalent to a homogeneous surface tension, which we assume can be absorbed in the surface tension term in Eq. 3 in the main text. The contribution in the second sum of Eq. (47) can be rewritten using the equilibrium concentration (46):

$$\begin{aligned}
\delta \mathcal{F}_{IJ} = & \int_{\mathcal{S}_I} dS_I \int_{\mathcal{S}_J} dS_J k_{\mathrm{B}} T c_0^2 \exp\left[-\frac{\phi\left(|\boldsymbol{X}_I - \boldsymbol{X}_J|\right)}{k_{\mathrm{B}}T}\right] \Big[ -\frac{1}{2}\left(g_I^{ij} \delta g_{I,ij} + g_J^{ij} \delta g_{J,ij}\right) \\
& + \phi'\left(|\boldsymbol{X}_I - \boldsymbol{X}_J|\right) \frac{X_I^\alpha - X_J^\alpha}{|\boldsymbol{X}_I - \boldsymbol{X}_J|} (\delta X_I^\alpha - \delta X_J^\alpha) \Big] \\
= & \ \delta \left\{ -\int_{\mathcal{S}_I} dS_I \int_{\mathcal{S}_J} dS_J k_{\mathrm{B}} T c_0^2 \exp\left[-\frac{\phi\left(|\boldsymbol{X}_I - \boldsymbol{X}_J|\right)}{k_{\mathrm{B}}T}\right] \right\} \ .
\end{aligned} \tag{49}$$

Thus, we can write an effective free energy

$$\mathcal{F}_{IJ} = \int_{\mathcal{S}_I} dS_I \int_{\mathcal{S}_J} dS_J \varphi_{IJ}\left(|\boldsymbol{X}_I - \boldsymbol{X}_J|\right) \ , \tag{50}$$

with

$$\varphi_{IJ} = -k_{\mathrm{B}} T c_0^2 \exp\left[-\frac{\phi\left(|\boldsymbol{X}_I - \boldsymbol{X}_J|\right)}{k_{\mathrm{B}}T}\right] \ . \tag{51}$$

In the main text we consider instead an effective Morse potential of the form

$$\varphi_{IJ} = D\left\{\left[1 - \exp\left(\frac{r_{\min} - |\boldsymbol{X}_I - \boldsymbol{X}_J|}{l}\right)\right]^2 - 1\right\} \ , \tag{52}$$

which like Eq. (51), has a minimum $-D$ at the position $|\boldsymbol{X}_I - \boldsymbol{X}_J| = r_{\min}$. The effective potential defined in Eq. (52) also has a strong repulsive component at short separation distance for $l \ll r_{\min}$.

Variations of Eq. (50) lead to Eq. (18) in the main text, using Eq. (14). In addition, we also show below that variation of the effective free energy $\mathcal{F}_{IJ}$ with respect to the surface shapes $I$, $J$, only depends on normal infinitesimal displacement of the surfaces, and so only depends on the surface shapes:

$$\delta \mathcal{F}_{IJ} = \int_{\mathcal{S}_I} dS_I \int_{\mathcal{S}_J} dS_J \Big\{ \varphi'\left(|\boldsymbol{X}_I - \boldsymbol{X}_J|\right) \frac{X_I^\alpha - X_J^\alpha}{|\boldsymbol{X}_I - \boldsymbol{X}_J|} (\delta X_I^\alpha - \delta X_J^\alpha)$$

$$
\begin{aligned}
&\quad + \varphi\left(|\boldsymbol{X}_I - \boldsymbol{X}_J|\right)\left(g_I^{ij}\left(\nabla_i \delta X_{Ij} + C_{Iij}\delta X_{In}\right) + g_J^{ij}\left(\nabla_i \delta X_{Jj} + C_{Jij}\delta X_{Jn}\right)\right\}, \\
&= \int_{\mathcal{S}_I} dS_I \int_{\mathcal{S}_J} dS_J \Big\{ \left[\partial_{Ii}\varphi\left(|\boldsymbol{X}_I - \boldsymbol{X}_J|\right)\right]\delta X_I^i + \left[\partial_{Ji}\varphi\left(|\boldsymbol{X}_I - \boldsymbol{X}_J|\right)\right]\delta X_J^i \\
&\quad + \varphi'\left(|\boldsymbol{X}_I - \boldsymbol{X}_J|\right)\frac{X_I^\alpha - X_J^\alpha}{|\boldsymbol{X}_I - \boldsymbol{X}_J|}\left(n_I^\alpha \delta X_{In} - n_J^\alpha \delta X_{Jn}\right) \\
&\quad - \left[\partial_{Ii}\varphi\left(|\boldsymbol{X}_I - \boldsymbol{X}_J|\right)\right]\delta X_I^i - \left[\partial_{Ji}\varphi\left(|\boldsymbol{X}_I - \boldsymbol{X}_J|\right)\right]\delta X_J^i \\
&\quad + \varphi\left(|\boldsymbol{X}_I - \boldsymbol{X}_J|\right)\left(C_{Ii}{}^i\delta X_{In} + C_{Ji}{}^i\delta X_{Jn}\right)\Big\} \\
&= \int_{\mathcal{S}_I} dS_I \int_{\mathcal{S}_J} dS_J \Big\{ \varphi'\left(|\boldsymbol{X}_I - \boldsymbol{X}_J|\right)\frac{X_I^\alpha - X_J^\alpha}{|\boldsymbol{X}_I - \boldsymbol{X}_J|}\left(n_I^\alpha \delta X_{In} - n_J^\alpha \delta X_{Jn}\right) \\
&\quad + \varphi\left(|\boldsymbol{X}_I - \boldsymbol{X}_J|\right)\left(C_{Ii}{}^i\delta X_{In} + C_{Ji}{}^i\delta X_{Jn}\right)\Big\},
\end{aligned} \tag{53}
$$

where we have used that:

$$
\partial_{Ii}\varphi\left(|\boldsymbol{X}_I - \boldsymbol{X}_J|\right) = \varphi'\left(|\boldsymbol{X}_I - \boldsymbol{X}_J|\right)\frac{\boldsymbol{X}_I - \boldsymbol{X}_J}{|\boldsymbol{X}_I - \boldsymbol{X}_J|} \cdot \boldsymbol{e}_{Ii}\ , \tag{54}
$$

and we generalise notations of differential geometry (S1 Appendix 1) to surfaces $I, J, ...$ by indicating with a $I, J, ...$ subscript the surface considered. The variation $\delta\mathcal{F}_{IJ}$ in Eq. (53) only depends on $\delta X_{In}$ and $\delta X_{Jn}$, which shows that the net tangential force (accounting for both the internal tension and the external force density) from the interaction potential is zero.

### 6 Discretised finite element equations

We discuss here the form of the finite element equations that we solve numerically. We consider a parameterisation of the shape $\mathcal{S}$ given in Eq. (8) in the main text. For the discretised surface,

$$
\delta \boldsymbol{X} = \sum_a \delta \boldsymbol{X}_a B_a\ , \tag{55}
$$

where notations are defined as in Eq.(8) in the main text. Using this decomposition and Eqs. (11), (17), and (18) for the variations of the metric and curvature tensors and Christoffel symbols, we have

$$
\delta g_{ij} = \sum_a \frac{\partial g_{ij}}{\partial \boldsymbol{X}_a} \cdot \delta \boldsymbol{X}_a\ , \tag{56}
$$

$$
\delta C_{ij} = \sum_a \frac{\partial C_{ij}}{\partial \boldsymbol{X}_a} \cdot \delta \boldsymbol{X}_a\ , \tag{57}
$$

$$
\delta \Gamma_{ij}^k = \sum_a \frac{\partial \Gamma_{ij}^k}{\partial \boldsymbol{X}_a} \cdot \delta \boldsymbol{X}_a\ , \tag{58}
$$

where we have defined

$$
\frac{\partial g_{ij}}{\partial \boldsymbol{X}_a} = (\partial_i B_a)\boldsymbol{e}_j + (\partial_j B_a)\boldsymbol{e}_i\ , \tag{59}
$$

$$
\frac{\partial C_{ij}}{\partial \boldsymbol{X}_a} = -\partial_i \boldsymbol{e}_j \cdot \frac{\partial \boldsymbol{n}}{\partial \boldsymbol{X}_a} - \boldsymbol{n} \cdot \partial_i \partial_j B_a\ , \tag{60}
$$

$$
\frac{\partial \Gamma_{ij}^k}{\partial \boldsymbol{X}_a} = g^{kl}\Gamma_{ij}^m \frac{\partial g_{lm}}{\partial \boldsymbol{X}_a} + g^{kl}(\partial_i \boldsymbol{e}_j)(\partial_l B_a) + \boldsymbol{e}^k \partial_i \partial_j B_a\ , \tag{61}
$$

and here

$$
\frac{\partial \boldsymbol{n}}{\partial \boldsymbol{X}_a} = -(\partial_i B_a)\boldsymbol{e}^i \otimes \boldsymbol{n}\ . \tag{62}
$$

#### 6.1 Active viscous layer with bending rigidity

We start by looking at the discretisation of an active viscous layer with bending rigidity. To obtain the equations governing the dynamics of the nodes of the mesh, we substitute Eq. (12) into Eq. (2) from the main text. Using the expressions in Eqs. (56) and (57), we get

$$
\delta W = \sum_a \boldsymbol{F}_a\left(\left\{\boldsymbol{X}_b^{(n)}, \boldsymbol{X}_b^{(n+1)}\right\}_{b\in\langle\langle a\rangle\rangle}, P^{(n+1)}\right) \cdot \delta \boldsymbol{X}_a = 0 \rightarrow \boldsymbol{F}_a\left(\left\{\boldsymbol{X}_b^{(n)}, \boldsymbol{X}_b^{(n+1)}\right\}_{b\in\langle\langle a\rangle\rangle}, P^{(n+1)}\right) = \boldsymbol{0}\ . \tag{63}
$$

Note that because of the compact nature of the basis functions $B_a$, the force $\boldsymbol{F}_a$ only depend on nodes that have non-zero basis functions in the neighbouring elements of node $a$, which are those formed by the first, second, and third nearest neighbours in the mesh (those connected to node $a$ by a 3-edge, 2-edge, or 1-edge path in the mesh) and we denote by $\langle\langle a \rangle\rangle$. As it will be apparent in the next section, these equations also depend on $P^{(n+1)}$, the intracellular pressure difference. We split the different terms leading to $\boldsymbol{F}_a$ individually for clarity:

$$\boldsymbol{F}_a = \boldsymbol{F}_{a,\text{friction}} + \boldsymbol{F}_{a,\text{viscous}} + \boldsymbol{F}_{a,\text{active}} + \boldsymbol{F}_{a,\text{Helfrich}} + \boldsymbol{F}_{a,\text{volume}} \;, \tag{64}$$

These terms are

$$\boldsymbol{F}_{a,\text{friction}} = \int_{\mathcal{S}^{(n)}} dS^{(n)}\; \xi \left( \frac{\boldsymbol{X}^{(n+1)} - \boldsymbol{X}^{(n)}}{\Delta t^{(n)}} \right) B_a \;, \tag{65}$$

$$\boldsymbol{F}_{a,\text{viscous}} = \int_{\mathcal{S}^{(n)}} dS^{(n)}\; \frac{1}{2}\eta g^{(n)ki} g^{(n)lj} \left( \frac{g_{kl}^{(n+1)} - g_{kl}^{(n)}}{\Delta t^{(n)}} \right) \frac{\partial g_{ij}^{(n+1)}}{\partial \boldsymbol{X}_a^{(n+1)}} \;, \tag{66}$$

$$\boldsymbol{F}_{a,\text{active}} = \int_{\mathcal{S}^{(n+1)}} dS^{(n+1)}\; \frac{1}{2}\gamma g^{(n+1),ij} \frac{\partial g_{ij}^{(n+1)}}{\partial \boldsymbol{X}_a^{(n+1)}} \;, \tag{67}$$

$$\begin{aligned}\boldsymbol{F}_{a,\text{Helfrich}} = \int_{\mathcal{S}^{(n+1)}} dS^{(n+1)}\; &\Bigg\{ \frac{\kappa}{4}\left( C^{(n+1)}{}_k^k - C_0 \right) \left[ \left( C^{(n+1)}{}_k^k - C_0 \right) g^{(n+1)ij} - 4C^{(n+1)ij} \right] \frac{\partial g_{ij}^{(n+1)}}{\partial \boldsymbol{X}_a^{(n+1)}} \\ &+ \kappa \left( C^{(n+1)}{}_k^k - C_0 \right) g^{(n+1)ij} \frac{\partial C_{ij}^{(n+1)}}{\partial \boldsymbol{X}_a^{(n+1)}} \Bigg\} \;.\end{aligned} \tag{68}$$

We note here that we integrate the dissipative terms Eqs. (65) and (66) on $\mathcal{S}^{(n)}$, whereas Eqs. (67) and (68) are integrated on $\mathcal{S}^{(n+1)}$; this guarantees that, when $\gamma$ is homogeneous and time-independent, the resulting time-integration scheme follows a gradient flow where the energy $\mathcal{F} = \int_{\mathcal{S}} (\gamma + \kappa(C_k{}^k - C_0)^2/2) dS$ decreases over time, which endows the discretisation with stability regardless of the value of $\Delta t^{(n)}$, see [3]. The term $\boldsymbol{F}_{a,\text{volume}}$ is discussed in the next section. To calculate the integrals in Eqs. (65)-(68), and since our surface is parametrised locally by finite element parametrisations of the form of Eq. (9) in the main text, we split integrals into a sum over integrals in each element. Integrals over elements are then transformed into integrals in the parametric domain of barycentric coordinates, i.e. given an integrand $f(\boldsymbol{X})$:

$$\int_{\mathcal{S}} dS f(\boldsymbol{X}) = \sum_e \int_{\mathcal{S}_e} dS f(\boldsymbol{X}) = \sum_e \int_0^1 ds_e^1 \int_0^{1-s^1} ds_e^2 f(\boldsymbol{X}(s_e^1, s_e^2)) \sqrt{g}(s_e^1, s_e^2) \;, \tag{69}$$

where $s_e^1$, $s_e^2$ denote the barycentric coordinates of element $e$. Using Gaussian integration, we approximate this integral by

$$\int_{\mathcal{S}} dS f = \sum_e \sum_{k=1}^{N_g} w_k f(\boldsymbol{X}(s_{ek}^1, s_{ek}^2, t)) \sqrt{g}(s_{ek}^1, s_{ek}^2) \;, \tag{70}$$

where $(s_{ek}^1, s_{ek}^2)$ is the parametric position of Gauss point $k$ and $w_k$ its associated weight, on element $e$. In practice, we loop over elements $e$ and over their Gauss points $k$, and calculate the partial sums to the expressions Eq. (65)-(68), which are then assembled following a typical finite element routine. To solve Eq. (63), which is a non-linear equation of the degrees of freedom $\boldsymbol{X}_a^{(n+1)}$, we use a Newton-Raphson method, which requires to solve Eq. (15) of the main text. For this method, we need to compute the matrix $\partial \boldsymbol{F}_a / \partial \boldsymbol{X}_b^{(n+1)}$. To obtain this matrix, we split the expressions as in Eq. (65)-(68),

$$\frac{\partial \boldsymbol{F}_{a,\text{friction}}}{\partial \boldsymbol{X}_b^{(n+1)}} = \frac{1}{\Delta t^{(n)}} \int_{\mathcal{S}^{(n)}} dS^{(n)}\; \xi B_a B_b \boldsymbol{I} \;, \tag{71}$$

$$\begin{aligned}\frac{\partial \boldsymbol{F}_{a,\text{viscous}}}{\partial \boldsymbol{X}_b^{(n+1)}} = \int_{\mathcal{S}^{(n)}} dS^{(n)}\; &\frac{1}{2}\eta g^{(n)ki} g^{(n)lj} \\ &\times \left[ \left( \frac{g_{kl}^{(n+1)} - g_{kl}^{(n)}}{\Delta t^{(n)}} \right) \frac{\partial^2 g_{ij}^{(n+1)}}{\partial \boldsymbol{X}_a^{(n+1)} \partial \boldsymbol{X}_b^{(n+1)}} + \frac{1}{\Delta t^{(n)}} \frac{\partial g_{ij}^{(n+1)}}{\partial \boldsymbol{X}_a^{(n+1)}} \otimes \frac{\partial g_{kl}^{(n+1)}}{\partial \boldsymbol{X}_b^{(n+1)}} \right] \;,\end{aligned} \tag{72}$$

$$\begin{aligned}\frac{\partial \boldsymbol{F}_{a,\text{active}}}{\partial \boldsymbol{X}_b^{(n+1)}} = \int_{\mathcal{S}^{(n+1)}} dS^{(n+1)}\; \frac{1}{2}\gamma &\left[ g^{(n+1)ij} \frac{\partial^2 g_{ij}^{(n+1)}}{\partial \boldsymbol{X}_a^{(n+1)} \partial \boldsymbol{X}_b^{(n+1)}} - g^{(n+1)ki} g^{(n+1)lj} \frac{\partial g_{ij}^{(n+1)}}{\partial \boldsymbol{X}_a^{(n+1)}} \otimes \frac{\partial g_{kl}^{(n+1)}}{\partial \boldsymbol{X}_b^{(n+1)}} \right. \\ &\left. + \frac{1}{2} g^{(n+1)ij} g^{(n+1)kl} \frac{\partial g_{ij}^{(n+1)}}{\partial \boldsymbol{X}_a^{(n+1)}} \otimes \frac{\partial g_{kl}^{(n+1)}}{\partial \boldsymbol{X}_b^{(n+1)}} \right] \;,\end{aligned} \tag{73}$$

$$
\begin{aligned}
\frac{\partial \boldsymbol{F}_{a,\mathrm{Helfrich}}}{\partial \boldsymbol{X}_b^{(n+1)}} = \int_{\mathcal{S}^{(n+1)}} dS^{(n+1)}\ \kappa \Bigg\{ & \frac{1}{4}\left(C^{(n+1)k}_{\ \ k} - C_0\right)\left[\left(C^{(n+1)k}_{\ \ k} - C_0\right) g^{(n+1)ij} - 4C^{(n+1)ij}\right] \frac{\partial^2 g_{ij}^{(n+1)}}{\partial \boldsymbol{X}_a^{(n+1)} \partial \boldsymbol{X}_b^{(n+1)}} \\
& + \Bigg\{ \left(C^{(n+1)k}_{\ \ k} - C_0\right)\Bigg[\frac{1}{4}\left(C^{(n+1)k}_{\ \ k} - C_0\right)\left(\frac{1}{2} g^{(n+1)ij} g^{(n+1)kl} - g^{(n+1)ik} g^{(n+1)jl}\right) \\
& \quad + g^{(n+1)ik} C^{(n+1)jl} + g^{(n+1)jl} C^{(n+1)ik} - \frac{1}{2} g^{(n+1)ij} C^{(n+1)kl} - \frac{1}{2} C^{(n+1)ij} g^{(n+1)kl}\Bigg] \\
& \quad + C^{(n+1)ij} C^{(n+1)kl}\Bigg\} \frac{\partial g_{ij}^{(n+1)}}{\partial \boldsymbol{X}_a^{(n+1)}} \otimes \frac{\partial g_{kl}^{(n+1)}}{\partial \boldsymbol{X}_b^{(n+1)}} \qquad (74) \\
& + \left(C^k_k - C_0\right) g_{ij}^{(n+1)} \frac{\partial^2 C_{ij}^{(n+1)}}{\partial \boldsymbol{X}_a^{(n+1)} \partial \boldsymbol{X}_b^{(n+1)}} + g^{(n+1)ij} g^{(n+1)kl} \frac{\partial C_{ij}^{(n+1)}}{\partial \boldsymbol{X}_a^{(n+1)}} \otimes \frac{\partial C_{kl}^{(n+1)}}{\partial \boldsymbol{X}_b^{(n+1)}} \\
& + \frac{1}{2}\left[\left(C^{(n+1)k}_{\ \ k} - C_0\right)\left(g^{(n+1)ij} g^{(n+1)kl} - 2 g^{(n+1)ik} g^{(n+1)jl}\right) - 2C^{(n+1)ij} g^{(n+1)kl}\right] \\
& \quad \times \left[\frac{\partial g_{ij}^{(n+1)}}{\partial \boldsymbol{X}_a} \otimes \frac{\partial C_{kl}^{(n+1)}}{\partial \boldsymbol{X}_b} + \frac{\partial C_{kl}^{(n+1)}}{\partial \boldsymbol{X}_a} \otimes \frac{\partial g_{ij}^{(n+1)}}{\partial \boldsymbol{X}_b}\right]\Bigg\}\ ,
\end{aligned}
$$

where $\boldsymbol{I}$ is the identity matrix in three dimensions, and

$$
\frac{\partial^2 g_{ij}}{\partial \boldsymbol{X}_a \partial \boldsymbol{X}_b} = \left[(\partial_i B_a)(\partial_j B_b) + (\partial_j B_a)(\partial_i B_b)\right] \boldsymbol{I}\ , \qquad (75)
$$

$$
\frac{\partial^2 C_{ij}}{\partial \boldsymbol{X}_a \partial \boldsymbol{X}_b} = -\partial_i \boldsymbol{e}_j \cdot \frac{\partial^2 \boldsymbol{n}}{\partial \boldsymbol{X}_a \partial \boldsymbol{X}_b} - \frac{\partial \boldsymbol{n}}{\partial \boldsymbol{X}_a} \partial_i \partial_j B_b - \frac{\partial \boldsymbol{n}}{\partial \boldsymbol{X}_b} \partial_i \partial_j B_a\ , \qquad (76)
$$

where

$$
\begin{aligned}
\frac{\partial^2 n^\alpha}{\partial X_a^\beta \partial X_b^\gamma} &= -\partial_i B_a \left(g^{ij} (\partial_j B_b) \delta^{\alpha\gamma} n^\beta - g^{ik} \frac{\partial g_{kj}}{\partial X_b^\gamma} e^{j\alpha} n^\beta + e^{i\alpha} \frac{\partial n^\beta}{\partial X_b^\gamma}\right) \\
&= -\partial_i B_a \left(g^{ij} (\partial_j B_b) \delta^{\alpha\gamma} n^\beta - g^{ik} ((\partial_k B_b) e_j^\gamma + (\partial_j B_b) e_k^\gamma) e^{j\alpha} n^\beta - e^{i\alpha} (\partial_j B_b) e^{j\beta} n^\gamma\right) \qquad (77) \\
&= (\partial_i B_a)(\partial_j B_b) \left[-g^{ij} \delta^{\alpha\gamma} n^\beta + g^{ij} e_k^\gamma e^{k\alpha} n^\beta + e^{i\gamma} e^{j\alpha} n^\beta + e^{i\alpha} e^{j\beta} n^\gamma\right] \\
&= (\partial_i B_a)(\partial_j B_b) \left[e^{i\alpha} e^{j\beta} n^\gamma + e^{j\alpha} n^\beta e^{i\gamma} - g^{ij} n^\alpha n^\gamma n^\beta\right] .
\end{aligned}
$$

#### 6.2 Adding cell pressure as a Lagrange multiplier

To add the cell pressure $P$ imposing the constraint in volume, we first note that the volume enclosed by $\mathcal{S}$ can be computed as a surface integral by applying the divergence theorem,

$$
V = \int_{\mathcal{V}} dV = \frac{1}{3} \int_{\mathcal{V}} dV\ \partial_\alpha X_\alpha = \frac{1}{3} \int_{\mathcal{S}} dS\ \boldsymbol{X} \cdot \boldsymbol{n}\ , \qquad (78)
$$

where $\mathcal{V}$ denotes the volume enclosed by $\mathcal{S}$. We note that the variation of volume can be written, by application of Reynold's transport theorem, as

$$
\delta V = \int_{\mathcal{S}} \delta \boldsymbol{X} \cdot \boldsymbol{n} dS\ . \qquad (79)
$$

Thus, by plugging the definition of the pressure force, Eq. (6) in the main text, into the principle of virtual work, Eq. (2) in the main text, we can write

$$
-P \int_{\mathcal{S}} \delta \boldsymbol{X} \cdot \boldsymbol{n} dS = -P \delta V\ . \qquad (80)
$$

The right hand side can then be interpreted as the second term in the variation of the Lagrangian $-\delta(P(V - V_0)) = -\delta P(V - V_0) - P\delta V$, with $V_0$ the cell target volume. This allows us to enforce the constraint $V^{(n+1)} = V_0$ through $P = P^{(n+1)}$. After discretisation, the right hand side of Eq. (80) leads to the following contribution to Eq. (64):

$$
\begin{aligned}
\boldsymbol{F}_{a,\mathrm{volume}} = -\frac{P^{(n+1)}}{3} \int_{\mathcal{S}^{(n+1)}} dS^{(n+1)} & \Bigg[B_a \boldsymbol{n}^{(n+1)} + \boldsymbol{X}^{(n+1)} \cdot \frac{\partial \boldsymbol{n}^{(n+1)}}{\partial \boldsymbol{X}_a^{(n+1)}} \\
& + \frac{1}{2} (\boldsymbol{X}^{(n+1)} \cdot \boldsymbol{n}^{(n+1)}) g^{(n+1)ij} \frac{\partial g_{ij}^{(n+1)}}{\partial \boldsymbol{X}_a^{(n+1)}}\Bigg] .
\end{aligned} \qquad (81)
$$

The constraint of prescribed volume can then be written as

$$F_P = \frac{1}{3}\int_{\mathcal{S}^{(n+1)}} dS^{(n+1)}\ \boldsymbol{X}^{(n+1)} \cdot \boldsymbol{n}^{(n+1)} = V_0\ , \tag{82}$$

which adds an extra equation to be solved. For the Newton-Raphson method, we need to compute the derivatives

$$\begin{aligned}
\frac{\partial \boldsymbol{F}_{a,\text{volume}}}{\partial \boldsymbol{X}_b^{(n+1)}} = -\frac{P^{(n+1)}}{3}\int_{\mathcal{S}^{(n+1)}} dS^{(n+1)} & \Bigg[ B_a \frac{\partial \boldsymbol{n}^{(n+1)}}{\partial \boldsymbol{X}_b^{(n+1)}} + \frac{\partial \boldsymbol{n}^{(n+1)}}{\partial \boldsymbol{X}_a^{(n+1)}} B_b + \boldsymbol{X}^{(n+1)} \cdot \frac{\partial^2 \boldsymbol{n}^{(n+1)}}{\partial \boldsymbol{X}_a^{(n+1)} \partial \boldsymbol{X}_b^{(n+1)}} \\
& + \frac{1}{2}\left( B_a \boldsymbol{n}^{(n+1)} + \boldsymbol{X}^{(n+1)} \cdot \frac{\partial \boldsymbol{n}^{(n+1)}}{\partial \boldsymbol{X}_a^{(n+1)}} \right) \otimes \left( g^{(n+1)ij} \frac{\partial g_{ij}^{(n+1)}}{\partial \boldsymbol{X}_b^{(n+1)}} \right) \\
& + \frac{1}{2}\left( g^{(n+1)ij} \frac{\partial g_{ij}^{(n+1)}}{\partial \boldsymbol{X}_a^{(n+1)}} \right) \otimes \left( B_b \boldsymbol{n}^{(n+1)} + \boldsymbol{X}^{(n+1)} \cdot \frac{\partial \boldsymbol{n}^{(n+1)}}{\partial \boldsymbol{X}_b^{(n+1)}} \right) \\
& + \frac{1}{2}(\boldsymbol{X}^{(n+1)} \cdot \boldsymbol{n}^{(n+1)}) g^{(n+1)ij} \frac{\partial^2 g_{ij}^{(n+1)}}{\partial \boldsymbol{X}_a^{(n+1)} \boldsymbol{X}_b^{(n+1)}} \\
& - \frac{1}{2}(\boldsymbol{X}^{(n+1)} \cdot \boldsymbol{n}^{(n+1)}) g^{(n+1)ik} g^{(n+1)jl} \frac{\partial g_{ij}^{(n+1)}}{\partial \boldsymbol{X}_a^{(n+1)}} \otimes \frac{\partial g_{kl}^{(n+1)}}{\partial \boldsymbol{X}_b^{(n+1)}} \\
& + \frac{1}{4}(\boldsymbol{X}^{(n+1)} \cdot \boldsymbol{n}^{(n+1)}) g^{(n+1)ij} g^{(n+1)kl} \frac{\partial g_{ij}^{(n+1)}}{\partial \boldsymbol{X}_a^{(n+1)}} \otimes \frac{\partial g_{kl}^{(n+1)}}{\partial \boldsymbol{X}_b^{(n+1)}} \Bigg] ,
\end{aligned} \tag{83}$$

$$\frac{\partial \boldsymbol{F}_{a,\text{volume}}}{\partial P} = -\frac{\partial F_P}{\partial \boldsymbol{X}_a^{(n+1)}} = \frac{1}{P^{(n+1)}} \boldsymbol{F}_{a,\text{volume}}\ . \tag{84}$$

#### 6.3 Adding cell-cell interactions

We now consider cell-cell interactions. Here, one needs to consider a larger system of equations, taking into account vertex forces calculated in the previous sections, but also force stemming from cell-cell interaction. For cell $I$ and node $a$, the condition $\delta W = 0$ defined in Eq. (17) in the main text now gives

$$\boldsymbol{F}_{I,a}\left( \left\{ \boldsymbol{X}_{J,b}^{(n)}, \boldsymbol{X}_{J,b}^{(n+1)} \right\}_{J,b \in \langle\langle I,a \rangle\rangle}, P_I^{(n+1)} \right) = \boldsymbol{0}\ . \tag{85}$$

Here $\langle\langle I,a \rangle\rangle$ identifies the set of vertices (identified as the pair of labels $J$ for the cell considered and $b$ for the vertex considered) that are neighbours of vertex $a$ in cell $I$. These contain all vertices that are neighbours to $a$ in cell $I$ and also all vertices that interact with node $a$ through the interaction potential. The function $\boldsymbol{F}_{I,a}$ then has the following decomposition:

$$\boldsymbol{F}_{I,a} = \boldsymbol{F}_{I,a,\text{friction}} + \boldsymbol{F}_{I,a,\text{viscous}} + \boldsymbol{F}_{I,a,\text{active}} + \boldsymbol{F}_{I,a,\text{Helfrich}} + \boldsymbol{F}_{I,a,\text{volume}} + \boldsymbol{F}_{I,a,\text{adhesion}}\ . \tag{86}$$

The 5 first terms in the right-hand side follow definitions obtained in the previous subsections. The remaining term, the contributions from adhesion, reads, using Eq. (18) in the main text:

$$\begin{aligned}
\boldsymbol{F}_{I,a,\text{adhesion}} = \int_{\mathcal{S}_I^{(n+1)}} dS_I^{(n+1)} \int_{\mathcal{S}_J^{(n+1)}} dS_J^{(n+1)} & \Bigg\{ \varphi'\left( \left| \boldsymbol{X}_I^{(n+1)} - \boldsymbol{X}_J^{(n+1)} \right| \right) \frac{\boldsymbol{X}_I^{(n+1)} - \boldsymbol{X}_J^{(n+1)}}{\left| \boldsymbol{X}_I^{(n+1)} - \boldsymbol{X}_J^{(n+1)} \right|} B_{I,a} \\
& + \frac{1}{2}\varphi\left( \left| \boldsymbol{X}_I^{(n+1)} - \boldsymbol{X}_J^{(n+1)} \right| \right) g_I^{(n+1)ij} \frac{\partial g_{Iij}^{(n+1)}}{\partial \boldsymbol{X}_{I,a}^{(n+1)}} \Bigg\} .
\end{aligned} \tag{87}$$

For the derivative of this force, $\partial \boldsymbol{F}_{I,a,\text{adhesion}} / \partial \boldsymbol{X}_{J,b}^{(n+1)}$, we distinguish between the case $J \neq I$

$$\begin{aligned}
\frac{\partial \boldsymbol{F}_{I,a,\text{adhesion}}}{\partial \boldsymbol{X}_{J,b}^{(n+1)}} = \int_{\mathcal{S}_I^{(n+1)}} dS_I^{(n+1)} \int_{\mathcal{S}_J^{(n+1)}} dS_J^{(n+1)} & \Bigg\{ \left[ \varphi''\left( \left| \boldsymbol{X}_I^{(n+1)} - \boldsymbol{X}_J^{(n+1)} \right| \right) - \frac{\varphi'\left( \left| \boldsymbol{X}_I^{(n+1)} - \boldsymbol{X}_J^{(n+1)} \right| \right)}{\left| \boldsymbol{X}_I^{(n+1)} - \boldsymbol{X}_J^{(n+1)} \right|} \right] \\
& \times \left( \frac{\boldsymbol{X}_I^{(n+1)} - \boldsymbol{X}_J^{(n+1)}}{\left| \boldsymbol{X}_I^{(n+1)} - \boldsymbol{X}_J^{(n+1)} \right|} \otimes \frac{\boldsymbol{X}_J^{(n+1)} - \boldsymbol{X}_I^{(n+1)}}{\left| \boldsymbol{X}_I^{(n+1)} - \boldsymbol{X}_J^{(n+1)} \right|} \right) B_{I,a} B_{J,b}
\end{aligned}$$

$$
\begin{aligned}
&- \frac{\varphi'\left(\left|\boldsymbol{X}_I^{(n+1)} - \boldsymbol{X}_J^{(n+1)}\right|\right)}{\left|\boldsymbol{X}_I^{(n+1)} - \boldsymbol{X}_J^{(n+1)}\right|} B_{I,a} B_{J,b} \boldsymbol{I} \qquad (88)\\
&+ \frac{1}{2}\left[\varphi'\left(\left|\boldsymbol{X}_I^{(n+1)} - \boldsymbol{X}_J^{(n+1)}\right|\right) B_{I,a} \frac{\boldsymbol{X}_I^{(n+1)} - \boldsymbol{X}_J^{(n+1)}}{\left|\boldsymbol{X}_I^{(n+1)} - \boldsymbol{X}_J^{(n+1)}\right|}\right] \otimes \left[g_J^{(n+1)ij} \frac{\partial g_{Jij}^{(n+1)}}{\partial \boldsymbol{X}_{J,b}^{(n+1)}}\right]\\
&+ \frac{1}{2}\left[g_I^{(n+1)ij} \frac{\partial g_{Iij}^{(n+1)}}{\partial \boldsymbol{X}_{I,a}^{(n+1)}}\right] \otimes \left[\varphi'\left(\left|\boldsymbol{X}_I^{(n+1)} - \boldsymbol{X}_J^{(n+1)}\right|\right) B_{J,b} \frac{\boldsymbol{X}_J^{(n+1)} - \boldsymbol{X}_I^{(n+1)}}{\left|\boldsymbol{X}_J^{(n+1)} - \boldsymbol{X}_I^{(n+1)}\right|}\right]\\
&+ \frac{1}{4}\varphi\left(\left|\boldsymbol{X}_I^{(n+1)} - \boldsymbol{X}_J^{(n+1)}\right|\right)\left[g_I^{(n+1)ij} \frac{\partial g_{Iij}^{(n+1)}}{\partial \boldsymbol{X}_{I,a}^{(n+1)}}\right] \otimes \left[g_J^{(n+1)ij} \frac{\partial g_{Jij}^{(n+1)}}{\partial \boldsymbol{X}_{J,b}^{(n+1)}}\right]\Bigg\},
\end{aligned}
$$

and the case $I = J$,

$$
\begin{aligned}
\frac{\partial \boldsymbol{F}_{I,a,\mathrm{adhesion}}}{\partial \boldsymbol{X}_{I,b}^{(n+1)}} = &\int_{\mathcal{S}_I^{(n+1)}} dS_I^{(n+1)} \int_{\mathcal{S}_J^{(n+1)}} dS_J^{(n+1)} \Bigg\{\left[\varphi''\left(\left|\boldsymbol{X}_I^{(n+1)} - \boldsymbol{X}_J^{(n+1)}\right|\right) - \frac{\varphi'\left(\left|\boldsymbol{X}_I^{(n+1)} - \boldsymbol{X}_J^{(n+1)}\right|\right)}{\left|\boldsymbol{X}_I^{(n+1)} - \boldsymbol{X}_J^{(n+1)}\right|}\right]\\
&\times \left(\frac{\boldsymbol{X}_I^{(n+1)} - \boldsymbol{X}_J^{(n+1)}}{\left|\boldsymbol{X}_I^{(n+1)} - \boldsymbol{X}_J^{(n+1)}\right|} \otimes \frac{\boldsymbol{X}_I^{(n+1)} - \boldsymbol{X}_J^{(n+1)}}{\left|\boldsymbol{X}_I^{(n+1)} - \boldsymbol{X}_J^{(n+1)}\right|}\right) B_{I,a} B_{I,b}\\
&+ \frac{\varphi'\left(\left|\boldsymbol{X}_I^{(n+1)} - \boldsymbol{X}_J^{(n+1)}\right|\right)}{\left|\boldsymbol{X}_I^{(n+1)} - \boldsymbol{X}_J^{(n+1)}\right|} B_{I,a} B_{I,b} \boldsymbol{I} \qquad (89)\\
&+ \frac{1}{2}\left[\varphi'\left(\left|\boldsymbol{X}_I^{(n+1)} - \boldsymbol{X}_J^{(n+1)}\right|\right) B_{I,a} \frac{\boldsymbol{X}_I^{(n+1)} - \boldsymbol{X}_J^{(n+1)}}{\left|\boldsymbol{X}_I^{(n+1)} - \boldsymbol{X}_J^{(n+1)}\right|}\right] \otimes \left[g_I^{(n+1)ij} \frac{\partial g_{Iij}^{(n+1)}}{\partial \boldsymbol{X}_{I,b}^{(n+1)}}\right]\\
&+ \frac{1}{2}\left[g_I^{(n+1)ij} \frac{\partial g_{Iij}^{(n+1)}}{\partial \boldsymbol{X}_{I,a}}\right] \otimes \left[\varphi'\left(\left|\boldsymbol{X}_I^{(n+1)} - \boldsymbol{X}_J^{(n+1)}\right|\right) B_{I,b} \frac{\boldsymbol{X}_I^{(n+1)} - \boldsymbol{X}_J^{(n+1)}}{\left|\boldsymbol{X}_J^{(n+1)} - \boldsymbol{X}_I^{(n+1)}\right|}\right]\\
&+ \frac{1}{4}\varphi\left(\left|\boldsymbol{X}_I^{(n+1)} - \boldsymbol{X}_J^{(n+1)}\right|\right)\left[g_I^{(n+1)ij} \frac{\partial g_{Iij}^{(n+1)}}{\partial \boldsymbol{X}_{I,a}^{(n+1)}}\right] \otimes \left[g_I^{(n+1)ij} \frac{\partial g_{Iij}^{(n+1)}}{\partial \boldsymbol{X}_{I,b}^{(n+1)}}\right]\\
&+ \frac{1}{2}\varphi\left(\left|\boldsymbol{X}_I^{(n+1)} - \boldsymbol{X}_J^{(n+1)}\right|\right)\Bigg[g_I^{(n+1)ij} \frac{\partial^2 g_{Iij}^{(n+1)}}{\partial \boldsymbol{X}_{I,a}^{(n+1)} \partial \boldsymbol{X}_{I,b}^{(n+1)}}\\
&\qquad - g_I^{(n+1)ik} g_I^{(n+1)jl} \frac{\partial g_{Iij}^{(n+1)}}{\partial \boldsymbol{X}_{I,a}^{(n+1)}} \otimes \frac{\partial g_{Ikl}^{(n+1)}}{\partial \boldsymbol{X}_{I,b}^{(n+1)}}\Bigg]\Bigg\}.
\end{aligned}
$$

We note that, together with Eq. (85), for each cell $I$ we solve $\boldsymbol{F}_{I,P} = V_0$ with the expression in Eq. (82) to impose the volume contraint through $P_I^{(n+1)}$.

### 7 Analytical solution for the flow on a spherical active surface

Here we derive an analytical expression for the flow on a spherical viscous surface resulting from gradient of active tensions, in the absence external torques. Our analysis is similar to the derivation in Ref. [4].

We separate here the velocity into its in-plane $v_i$ and normal $v_n$ components. We further consider the Hodge decomposition of the tangential velocity [5, 6],

$$
v_i = \partial_i \phi + \epsilon_i{}^j \partial_j \psi \,, \qquad (90)
$$

where $\phi$ and $\psi$ are the irrotational and solenoidal potentials, i.e. $\partial_i \phi$ is an irrotational field whereas $\epsilon_i{}^j \partial_j \psi$ is a solenoidal field. In this decomposition, we assume the surface is simply-connected as otherwise there would be another, harmonic component of $v_i$ [5, 6]. Using the constitutive equations (12) in the main text, the tension tensor components are:

$$
t^{ij} = 2\eta v^{ij} + \gamma g^{ij} + \kappa (C_k{}^k - C_0)\left(\frac{1}{2}(C_k{}^k - C_0) g^{ij} - C^{ij}\right) \,, \qquad (91)
$$

$$t_n^i = \kappa \partial^i C_k{}^k \quad . \tag{92}$$

where we have considered that the bending modulus $\kappa$ and preferred curvature $C_0$ are uniform on the surface, while $\gamma$ can be inhomogeneous. The external force density is given by Eq. (5) in the main text, and arises from an external pressure and an effective external friction force density. The in-plane and normal force balance equations (24) and (25) can then be written as:

$$\eta \left[2\Delta\partial^i\phi + \left[\epsilon^{jk}\nabla_j\nabla^i + \epsilon^{ik}\Delta\right]\partial_k\psi + 2\nabla_j\left(C^{ij}v_n\right)\right] \\ -\xi\left[\partial^i\phi + \epsilon^{ij}\partial_j\psi\right] + \partial^i\gamma = 0 \ . \tag{93}$$

$$\kappa\left[\Delta C_k{}^k - (C_k{}^k - C_0)\left(\frac{1}{2}C_i{}^i(C_j{}^j - C_0) - C^{ij}C_{ij}\right)\right] \\ -2\eta\left\{C^{ij}\nabla_i\partial_j\phi + \epsilon_{ij}C^{ik}\nabla_k\partial^j\psi + C_{ij}C^{ij}v_n\right\} - \gamma C_i{}^i + P - \xi v_n = 0 \ , \tag{94}$$

where $\Delta = \nabla_i\nabla^i$.

We now introduce three arbitrary scalar fields $w_\phi$, $w_\psi$ and $w_n$. We multiply Eq. (93) by $\partial_i w_\phi$ and by $\epsilon_i{}^j\partial_j w_\psi$, and Eq. (94) by $w_n$ and integrate the resulting equations on $\mathcal{S}$. We obtain:

$$-\int_{\mathcal{S}} dS\left\{2\eta(\nabla_j\partial_i w_\phi)\left[\nabla^j\partial^i\phi + \epsilon^{ik}\nabla^j\partial_k\psi + C^{ij}v_n\right] + \Delta w_\phi\left[\gamma - \xi\phi\right]\right\} = 0 \ , \tag{95}$$

$$\int_{\mathcal{S}} dS\left\{-2\eta\left[\epsilon_i{}^l(\nabla_j\partial_l w_\psi)\nabla^j\partial^i\phi + (\nabla_j\partial_i w_\psi)\nabla^j\partial^i\psi - \frac{1}{2}(\Delta w_\psi)\Delta\psi \right.\right. \\ \left.\left. +\epsilon_i{}^l(\nabla_j\partial_l w_\psi)C^{ij}v_n\right] + \xi\psi\Delta w_\psi\right\} = 0 \ , \tag{96}$$

$$\int_{\mathcal{S}} dS w_n\left\{\kappa\left[\Delta C_k{}^k - (C_k{}^k - C_0)\left(\frac{1}{2}C_l{}^l(C_m{}^m - C_0) - C^{ij}C_{ij}\right)\right]\right. \\ \left. -2\eta\left\{C^{ij}\nabla_i\partial_j\phi + \epsilon_{ij}C^{ik}\nabla_k\partial^j\psi + C_{ij}C^{ij}v_n\right\} - \gamma C_i{}^i + P - \xi v_n\right\} = 0 \ , \tag{97}$$

where we have integrated by parts some of the terms. Taking into account that $\nabla_i\partial_j A = \nabla_j\partial_i A$ with $A$ a scalar and that $\nabla_i\nabla_j B_k = \nabla_j\nabla_i B_k - R^l{}_{kij}B_l$ with $\boldsymbol{B}$ a vector and

$$R_{lkij} = K\left(g_{li}g_{kj} - g_{lj}g_{ki}\right) \ , \tag{98}$$

the Riemann tensor on a surface of Gaussian curvature $K = \det(C_i^j)$, and assuming that $\mathcal{S}$ is a sphere of radius $\ell$ ($C_{ij} = g_{ij}/\ell$, $K = 1/\ell^2$), we obtain:

$$\int_{\mathcal{S}} dS\,\Delta w_\phi\left\{2\eta\left[\left(\frac{1}{\ell^2} + \Delta\right)\phi + \frac{v_n}{\ell}\right] + \gamma - \xi\phi\right\} = 0 \ , \tag{99}$$

$$\int_{\mathcal{S}} dS\,\Delta w_\psi\left\{\eta\left(\frac{2}{\ell^2} + \Delta\right)\psi - \xi\psi\right\} = 0 \ , \tag{100}$$

$$\int_{\mathcal{S}} dS\,w_n\left\{\kappa\left(\frac{2}{\ell} - C_0\right)\frac{C_0}{\ell} - \frac{2\eta}{\ell}\Delta\phi - \frac{2\gamma}{\ell} + P - \left(\frac{4\eta}{\ell^2} + \xi\right)v_n\right\} = 0 \ . \tag{101}$$

We now expand $\phi = \sum_A\sum_a Y^{Aa}\phi^{Aa}$, $\psi = \sum_A\sum_a Y^{Aa}\psi^{Aa}$, $v_n = \sum_A\sum_a Y^{Aa}v_n^{Aa}$, and $\gamma = \sum_A\sum_a Y^{Aa}\gamma^{Aa}$ with $Y^{Aa}$ the spherical harmonic of degree $A$ and order $a$, with $-A \leq a \leq A$. Defining spherical coordinates $\theta$, $\phi$, such that a point of the sphere of radius $\ell$ is given by

$$\boldsymbol{X}(\theta,\phi) = \ell(\sin\theta\cos\phi\tilde{\boldsymbol{e}}_x + \sin\theta\sin\phi\tilde{\boldsymbol{e}}_y + \cos\theta\tilde{\boldsymbol{e}}_z) \ , \tag{102}$$

the spherical harmonics are then given by

$$Y^{Aa}(\theta,\phi) = ZP^{Aa}(\cos\theta)e^{-ia\varphi} \ , \tag{103}$$

with $P^{Aa}$ the associated Legendre polynomial of degree $A$ and order $a$, and $Z$ a normalising constant such that $\int_{\mathcal{S}}|Y^{Aa}|^2 dS = 1$, satisfying

$$\Delta Y^{Aa} = -\frac{A(A+1)}{\ell^2}Y^{Aa} \ . \tag{104}$$

Imposing $w_\phi = w_\psi = w_n = Y^{Bb}$, and taking into account the orthonormality of spherical harmonics, i.e. $\int_{\mathcal{S}} Y^{Aa}Y^{Bb*}dS = \delta^{AB}\delta^{ab}$ with $^*$ indicating the complex conjugate, we obtain the following algebraic equations:

$$\left(A(A+1) - 1 + \frac{\xi\ell^2}{2\eta}\right)\phi^{Aa} - v_n^{Aa}\ell - \frac{\ell^2}{2\eta}\gamma^{Aa} = 0 \ , \quad A > 0 \ , \tag{105}$$

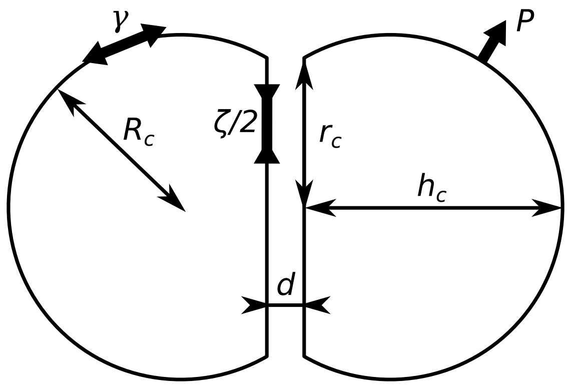


**Figure 1:** Diagram of a cell doublet as two mirror spherical caps of basal radius $r_c$, height $h_c$, radius of curvature $R_c$ and separated by a distance $d$. The active tension $\gamma$ is balanced by the pressure $P$ according to the law of Laplace. Linkers generate an effective adhesion tension $\zeta/2$ per cell.

$$\left(A(A+1) - 2 + \frac{\xi\ell^2}{\eta}\right)\psi^{Aa} = 0\ , \quad A > 0, \tag{106}$$

$$A(A+1)\phi^{Aa} - \frac{\ell^2}{\eta}\gamma^{Aa} + \frac{\sqrt{4\pi}\ell^3}{2\eta}\delta^{A0}\left[P\ell + \kappa\left(\frac{2}{\ell} - C_0\right)C_0\right] - \ell\left(2 + \frac{\xi\ell^2}{2\eta}\right)v_n^{Aa} = 0\ , \tag{107}$$

where Eq. (107) holds for all $A \geq 0$. We note that the zero-th mode $A = a = 0$ for $\phi$ and $\psi$ is arbitrary since uniform values of $\phi$ and $\psi$ do not change the velocity field defined by Eq. (90); we choose here that they vanish for simplicity. Given $\gamma^{Aa}$ and $P$, the solution is

$$\phi^{Aa} = \frac{\gamma^{Aa}\frac{\xi\ell^4}{4\eta^2}}{A(A+1)(1+\frac{\xi\ell^2}{2\eta}) + (2+\frac{\xi\ell^2}{2\eta})(\frac{\xi\ell^2}{2\eta} - 1)}, \quad A > 0\ , \tag{108}$$

$$\psi^{Aa} = 0, \qquad A > 0\ , \tag{109}$$

$$v_n^{Aa} = \frac{1}{\left(2+\frac{\xi\ell^2}{2\eta}\right)\ell}\left[A(A+1)\phi^{Aa} - \frac{\ell^2}{\eta}\gamma^{Aa} + \frac{\sqrt{4\pi}\ell^3}{2\eta}\delta^{A0}\left(P\ell + \kappa\left(\frac{2}{\ell} - C_0\right)C_0\right)\right]\ . \tag{110}$$

### 8 Approximate description of a cell doublet

Here we compute an approximate solution for the cell doublet valid in the limit of $\hat{r}_{\min}, \tilde{l} \ll 1$. We assume that the doublet is formed by two mirror spherical caps of height $h_c$ and base radius $r_c$ separated by a distance $d$, see Fig. 1. Since $\tilde{l} \ll 1$, only the two circular discs at the base of each cell interact. We label a point $\boldsymbol{X}_1$ on the contact disc of cell 1 by polar coordinates $r_1$, $\theta_1$, and we define a point on the contact disc of the second cell by:

$$\boldsymbol{X}_2 = \boldsymbol{X}_1 + r\cos\theta\tilde{\boldsymbol{e}}_x + r\sin\theta\tilde{\boldsymbol{e}}_y + d\tilde{\boldsymbol{e}}_z\ , \tag{111}$$

where the $z$ axis goes along the line joining the cell centers. The distance between two points on the two surfaces is then $|\boldsymbol{X}_2 - \boldsymbol{X}_1| = \sqrt{d^2 + r^2}$. The interaction energy can be written as:

$$\mathcal{F}_{IJ} = \int_0^{r_c} dr_1 \int_0^{2\pi} r_1 d\theta_1 \int_{\mathcal{S}_2(r_1,\theta_1)} r dr d\theta\, \varphi\left(\sqrt{d^2+r^2}\right)\ , \tag{112}$$

where the second integral is taken over the domain $\mathcal{S}_2(r_1, \theta_1)$ corresponding to points on cell 2 which are labelled by $r, \theta$ and are within the circular disc. Here we approximate this integral as an integral on the infinite plane; as the interaction potential $\varphi$ is short-ranged, this is valid sufficiently far away from the boundaries of the contact discs. Therefore, the approximation that applies when the contact disc is much larger than $l$, $r_{\min}$. One then obtains:

$$\mathcal{F}_{IJ} = \pi r_c^2 \left[2\pi \int_0^\infty dr\, r\varphi\left(\sqrt{r^2+d^2}\right)\right] \tag{113}$$

$$= \pi r_c^2 \zeta(d) \ , \tag{114}$$

where $\zeta(d) = 2\pi \int_0^\infty dr r \varphi\left(\sqrt{r^2+d^2}\right)$ plays the role of a negative surface tension of the adhesion patch; note that the tension per cell is thus $\zeta(d)/2$. In Eq. (112) we have used that $\hat{r}_{\min}, \tilde{l} \ll 1$ to approximate the integral over the circle of radius $r_c$ by an integral on an infinite plane. Adding this to the effective energy accounting for the homogeneous active tension $\bar{\gamma}$ and the pressure $P$, we have

$$\mathcal{F}(r_c, h_c, d, P) = 2\bar{\gamma} A + \pi r_c^2 \zeta(d) - 2P(V - V_0) \ . \tag{115}$$

where $A = \pi(2r_c^2 + h_c^2)$ is the area of each cell, $V = \pi h_c(3r_c^2 + h_c^2)/6$ is the cell volume, and $P$ the cell pressure. To find the equilibrium condition $(r_c^*, h_c^*, d^*, P^*)$, we take variations of this energy. Variations with respect to $d$ define $d^*$ through

$$\zeta'(d^*) = 0 \ . \tag{116}$$

Variations with respect to $h_c$ lead to law of Laplace

$$4\bar{\gamma} h_c - P(r_c^2 + h_c^2) = 0 \rightarrow P = \frac{2\bar{\gamma}}{R_c} \ , \tag{117}$$

where we have used that the radius of curvature of the cap is $R_c = (r_c^2 + h_c^2)/(2h_c)$. Together with Eq. (117), variations with respect to $r_c$ lead to the law of Young-Dupré

$$\begin{aligned} & 8\bar{\gamma} r_c + 2r_c \zeta(d^*) - 2P h_c r_c = 8\bar{\gamma} r_c + 2r_c \zeta(d^*) - 4\bar{\gamma} \frac{h_c r_c}{R_c} = 0 \\ & \rightarrow 2\bar{\gamma}\left(2 - \frac{h_c}{R_c}\right) + \zeta^* = 0 \ . \end{aligned} \tag{118}$$

where we introduced the notation $\zeta^* = \zeta(d^*)$. Finally, variations with respect to $P$ lead to the volume constraint

$$\frac{1}{6}\pi h_c(3r_c^2 + h_c^2) = V_0 \rightarrow r_c^* = \sqrt{\frac{2V_0}{\pi h_c^*} - \frac{1}{3}(h_c^*)^2} \ . \tag{119}$$

Substituting this expression in the definition of $R_c$ gives $R_c = V_0/(\pi h_c^2) + h_c/3$ and in Eq. (118),

$$2\bar{\gamma}\left(2 - \frac{h_c}{V_0/(\pi h_c^2) + h_c/3}\right) + \zeta^* = 0 \rightarrow h_c^* = \left(\frac{3V_0(4\bar{\gamma} + \zeta^*)}{\pi(2\bar{\gamma} - \zeta^*)}\right)^{1/3} \ . \tag{120}$$

For a Morse potential of the form of Eq. (52), one can analytically compute $\zeta(d)$,

$$\zeta_{\text{Morse}}(d) = D\pi\left[\frac{1}{2} l(l+2d) e^{-\frac{2(d-r_{\min})}{l}} - 4l(l+d)e^{\frac{r_{\min}-d}{l}}\right] \ , \tag{121}$$

whose minimum is at $d^* = r_{\min} - l \ln 2$. In the limit $l \ll r_{\min}$, one then has $\zeta^*_{\text{Morse}} = -\beta D l r_{\min}$ with $\beta = 4\pi$.

For the potential with a cutoff that we employ in our numerical simulations (Eq. (26) in the main text), the value of $\zeta^*$ can be computed numerically. Taking $r_1 = r_{\min}$ and $r_2 = r_{\min} + 3l$ and $r_{\min} = 3l$, one finds $\zeta^* = -\beta D r_{\min} l$ with $\beta \simeq 10.7$. The effective tension at the interface at equilibrium is then

$$2\bar{\gamma} + \zeta^* = 2\bar{\gamma} - \beta D r_{\min} l \ , \tag{122}$$

which becomes negative, and thus we expect to get a buckling instability, when

$$\frac{D r_{\min} l}{\bar{\gamma}} = \frac{2}{\beta} \ . \tag{123}$$

### 9 Approximate description of a planar, regular cell packing

Here we discuss approximate models for the shape of cells organised within a planar sheet, as in Fig. 5A in the main text. We consider that cells have a fixed volume $V_0$, a surface tension for free interfaces $\gamma^{ab}$, and a surface tension for interfaces in contact with other cells, $\gamma^l < \gamma^{ab}$. The effective energy of a single cell can then be written:

$$\mathcal{F} = 2\gamma^{ab} A^{ab} + \gamma^l A^l - P(V - V_0) \ , \tag{124}$$

where $A^{ab}$ is the apical surface area, equal to the basal surface area, $A^l$ is the lateral or contact surface area, $V$ the cell volume and $P$ the cellular pressure difference.

Treating each cell as an hexagonal prism with side length $a$ and height $h_c$, one has $A^{ab} = 3\sqrt{3}a^2/2$ the apical and basal surface area, $A^l = 6ah_c$ the lateral surface area, and the cell volume is $V = 3\sqrt{3}a^2h_c/2$. In that case:

$$\frac{2A^{ab}}{A^l} = \frac{\gamma^l}{2\gamma^{ab}} \ . \tag{125}$$

Alternatively, one can approximate the shape of a cell as a cylinder of radius $r_c$ and height $h_c$, connected to two spherical caps with height $h_s$. Then $A^{ab} = \pi(r_c^2 + h_s^2)$, $A^l = 2\pi r_c h_c$ and $V = \pi r_c^2 h_c + 2\pi h_s(3r_c^2 + h_s^2)/6$. Minimisation of $\mathcal{F}$ leads to:

$$\frac{2A^{ab}}{A^l} = \frac{\gamma^{ab}}{\gamma^l}\left(\frac{\gamma^{ab}}{\sqrt{\gamma^{ab2} - \gamma^{l2}}} - 1\right) \ . \tag{126}$$

One can also envision a "hybrid" model, where the cell is considered as a union of an hexagonal prism of height $h_c$ and side length $r_c$, and two spherical caps or height $h_s$ and radius $r_c$. We consider an approximate case, in that the extra surface area that arises from joining the prism hexagonal surfaces to the spherical caps circular discs, is neglected. In that case, one has $A^{ab} = \pi(r_c^2 + h_s^2)$, $A^l = 6r_c h_c$ and the cell volume is given by $V = 3\sqrt{3}r_c^2 h_c/2 + 2\pi h_s(3r_c^2 + h_s^2)/6$. Minimisation of $\mathcal{F}$ leads to:

$$\frac{2A^{ab}}{A^l} = \frac{3\gamma^{ab}\left(\frac{\sqrt{3}\gamma^{ab}\left(3\gamma^{ab2} - 4\gamma^{l2}\right)}{\sqrt{\left(3\gamma^{ab2} - 4\gamma^{l2}\right)^3}} - 1\right)}{4\gamma^l}. \tag{127}$$

The solution of these 3 models, together with results from simulations, are plotted in Fig. 5E. In all cases, one takes $\gamma^{ab} = \bar{\gamma}$ and

$$\gamma^l = \bar{\gamma}\left(1 - \frac{\beta\tilde{D}}{2}\right) \ , \tag{128}$$

which is the effective surface tension of an adhering interface, for one of the two participating cells, in the limit where the interface is large compared to $l$, $r_{min}$ (Eq. (122)).
