## Supplementary figures and images for "Interacting active surfaces: a model for three-dimensional cell aggregates"

### S2 Figure

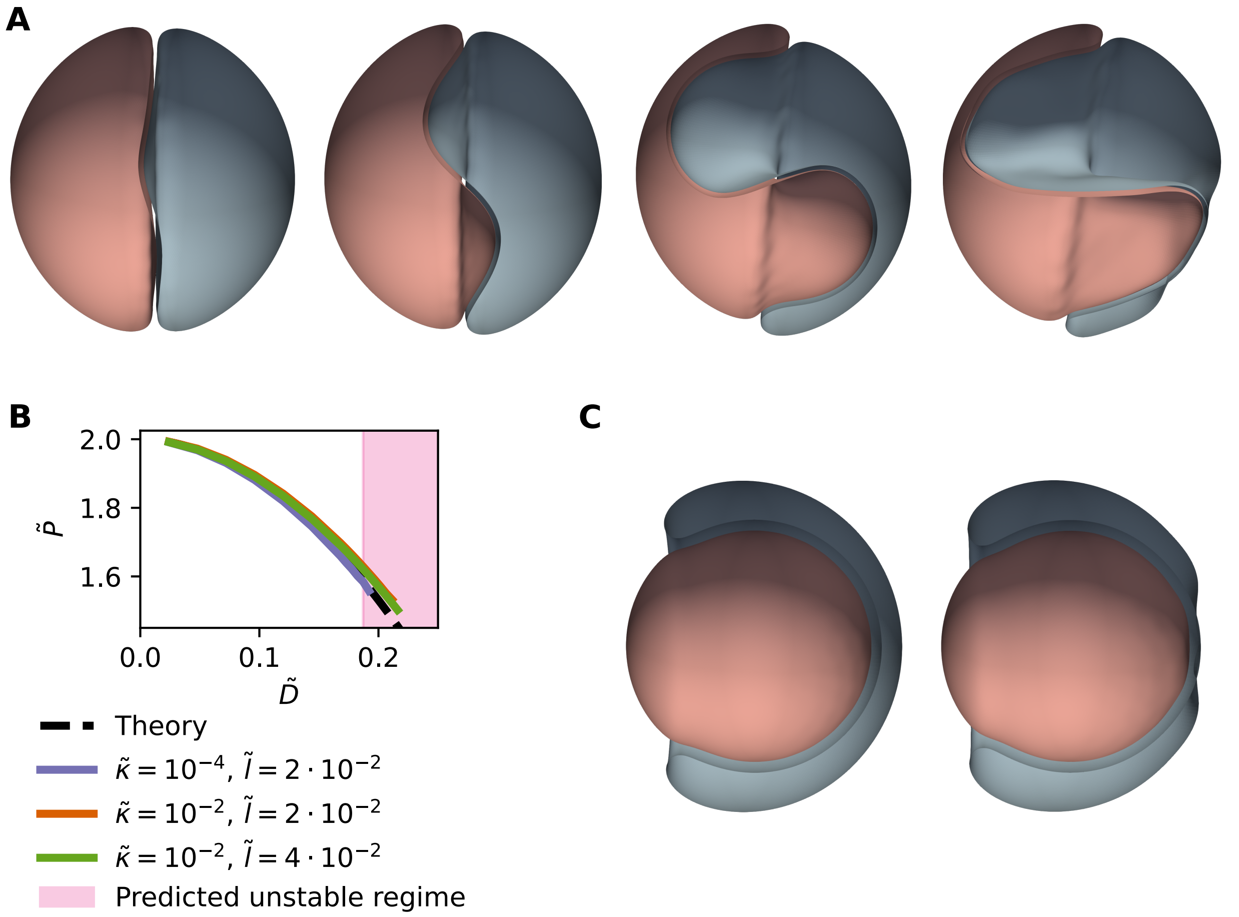
